# Multiplexed profiling of kinase interactomes quantifies cellular network plasticity

**DOI:** 10.1101/2021.09.14.460283

**Authors:** Martin Golkowski, Andrea Lius, Tanmay Sapre, Ho-Tak Lau, Taylor Moreno, Dustin J. Maly, Shao-En Ong

**Author notes:** Corresponding Authors (S-E.O.), (M.G.).

## Abstract

Cells utilize protein-protein interaction (PPI) networks to receive, transduce, and respond to stimuli. Interaction network rewiring drives devastating diseases like cancers, making PPIs attractive targets for pharmacological intervention. Kinases are druggable nodes in PPI networks but high-throughput proteomics approaches to quantify disease-associated kinome PPI rewiring are lacking. We introduce kinobead competition and correlation analysis (Ki-CCA), a chemoproteomics approach to simultaneously map hundreds of endogenous kinase PPIs. We identified 2,305 PPIs of 300 kinases across 18 diverse cancer lines, quantifying the high plasticity of interaction networks between cancer types, signaling, and phenotypic states; this database of dynamic kinome PPIs provides deep insights into cancer cell signaling. We discovered an AAK1 complex promoting epithelial-mesenchymal transition and drug resistance, and depleting its components sensitized cells to targeted therapy. Ki-CCA enables rapid and highly multiplexed mapping of kinome PPIs in native cell and tissue lysates, without epitope tagged baits, protein labeling, or antibodies.

## INTRODUCTION

Human diseases, including inflammatory and neurological disorders, infections, and cancers, dynamically alter cellular protein-protein interaction (PPI) networks in their natural progression as well as in response to therapeutic intervention ^1–4^. To understand disease mechanisms and to prioritize proteins for drug target discovery, it is therefore crucial to map PPI networks in a context-dependent manner. Protein kinases and phosphatases regulate cellular PPI networks through protein phosphorylation and de-phosphorylation, assembling and disassembling receptors, adapters, effectors, and their substrates into signaling complexes and molecular machines ^5^. Unsurprisingly, kinases are frequently dysregulated in human disease. Furthermore, kinases are highly druggable with synthetic ATP analogues, and are one of the most important classes of drug targets ^6,7^. Consequently, proteomics approaches that broadly map altered kinase PPI networks in specific disease contexts hold great promise to identify novel therapeutic targets ^8^.

Mass spectrometry (MS)-based interactomics approaches such as affinity purification-MS (AP-MS) and proximity labeling-MS strategies like BioID are powerful methods for large-scale mapping of PPI networks ^9^. Globally identifying PPIs in a single model cell line model using such approaches, however, requires the generation hundreds to thousands of stable cell lines that express epitope-tagged bait proteins, and a two- to three-fold larger number of affinity pulldowns and LC-MS experiments ^10,11^. For instance, Huttlin et al. mapped the cellular interactomes of two cell lines, HEK293T and HCT116, performing 15,650 affinity pulldowns that were analyzed in 31,300 LC-MS runs. This study yielded a comprehensive interactomics database of 170,280 interactions between 14,550 proteins, revealing that PPI network composition is highly context dependent. This study also showcases the limitations AP-MS, i.e., that comparing PPI network dynamics across disease models, cell signaling, and phenotypic states is low-throughput and extremely labor intensive. Furthermore, expressing exogenous epitope-tagged baits or BioID fusion proteins can alter the native interactome of proteins, potentially obscuring the biological function of endogenous proteins. The application of genetic fusions in clonal cell lines further complicates the comparison of assembled protein complexes across multiple cell states and conditions. Novel approaches such as size exclusion chromatography (SEC) coupled to a LC-MS readout tremendously increased interactomics throughput ^12–14^. Yet, SEC-MS has a lower dynamic range compared to AP-based methods, disfavoring the identification of low-abundance PPIs such as interactions between certain kinases and their signaling effectors.

To allow sensitive, rapid, and multiplexed PPI mapping of the kinome without the need for exogenous genetic tagging, bait expression, or antibodies, we introduce <u>ki</u>nobead <u>c</u>ompetition and <u>c</u>orrelation <u>a</u>nalysis (Ki-CCA). Kinobeads, also known as multiplexed inhibitor beads (MIBs) ^15,16^, enrich most expressed kinases along with thousands of interacting, non-kinase, proteins for sensitive identification and quantification by LC-MS ^17–19^. To allow high confidence, multiplexed identification of kinobead-bound kinase complexes, our chemoproteomics approach combines competition of kinases and their interaction partners using a set of 21 of broadly selective kinase inhibitor (KI) probes, LC-MS analysis, and Pearson correlation of MS intensity values to identify patterns of competed kinases complexes.

We used our Ki-CCA to globally profile kinase PPI networks in 18 diverse cancer cell lines and HeLa cells in different signaling states (21 conditions total) by performing 792 label-free kinobead/LC-MS soluble competition experiments. We found that our Ki-CCA identified and quantified 2,305 PPIs between 300 kinases and 1,436 proteins across 21 cell lines and cell states, with the number of interactions per kinase ranging between 1 and 103. Our kinome interactomics dataset presents an invaluable resource that gives deep insights into how cancer types, time-resolved signaling, and phenotypic states shape kinase interaction networks. For instance, analyzing an *AXL* RNAi knockdown model of cancer cell epithelial-mesenchymal transition (EMT), we discovered that cancer cell phenotypic plasticity is associated with a kinome-wide rewiring of kinase PPIs. Among EMT-associated PPIs we identified an adapter-associated kinase 1 (AAK1) complex that is highly abundant in drug resistant and mesenchymal-like hepatocellular carcinoma (HCC) cells compared to drug-sensitive epithelial-like HCC cells. RNAi knockdown of AAK1 complex components greatly affected EMT marker expression and sensitized cells to targeted therapy, demonstrating that Ki-CCA can identify kinases and kinase-interacting proteins that may serve as potential cancer drug targets.

## RESULTS

### Highly multiplexed profiling of native kinase interaction networks

To develop an approach that unbiasedly identifies and quantifies native kinase complexes present in cells and tissues, we exploited the kinobeads’ ability to enrich most expressed kinases and many of their interactors. We have previously shown that specific kinobead-bound kinase complexes can be identified by soluble competition with selective KIs (Figure 1A) ^19,20^. To simultaneously identify the hundreds of kinobead-bound kinase complexes, we envisioned an approach where a limited number broad-selectivity KIs with orthogonal kinase binding affinities, hereafter referred to as kinase interactome probes (KIPs), would compete for binding of most kinases and their interactors to the kinobeads, producing distinct enrichment profiles for each kinase’s interactome. Performing multiple competition experiments each using a different KIP and correlating MS intensity values of all competed kinases with all competed non-kinase proteins should then identify specific kinase-interactor pairs with high positive r-values (<u>ki</u>nobead <u>c</u>ompetition and <u>c</u>orrelation <u>a</u>nalysis, CCA, Figure 1B).

**Figure 1.**
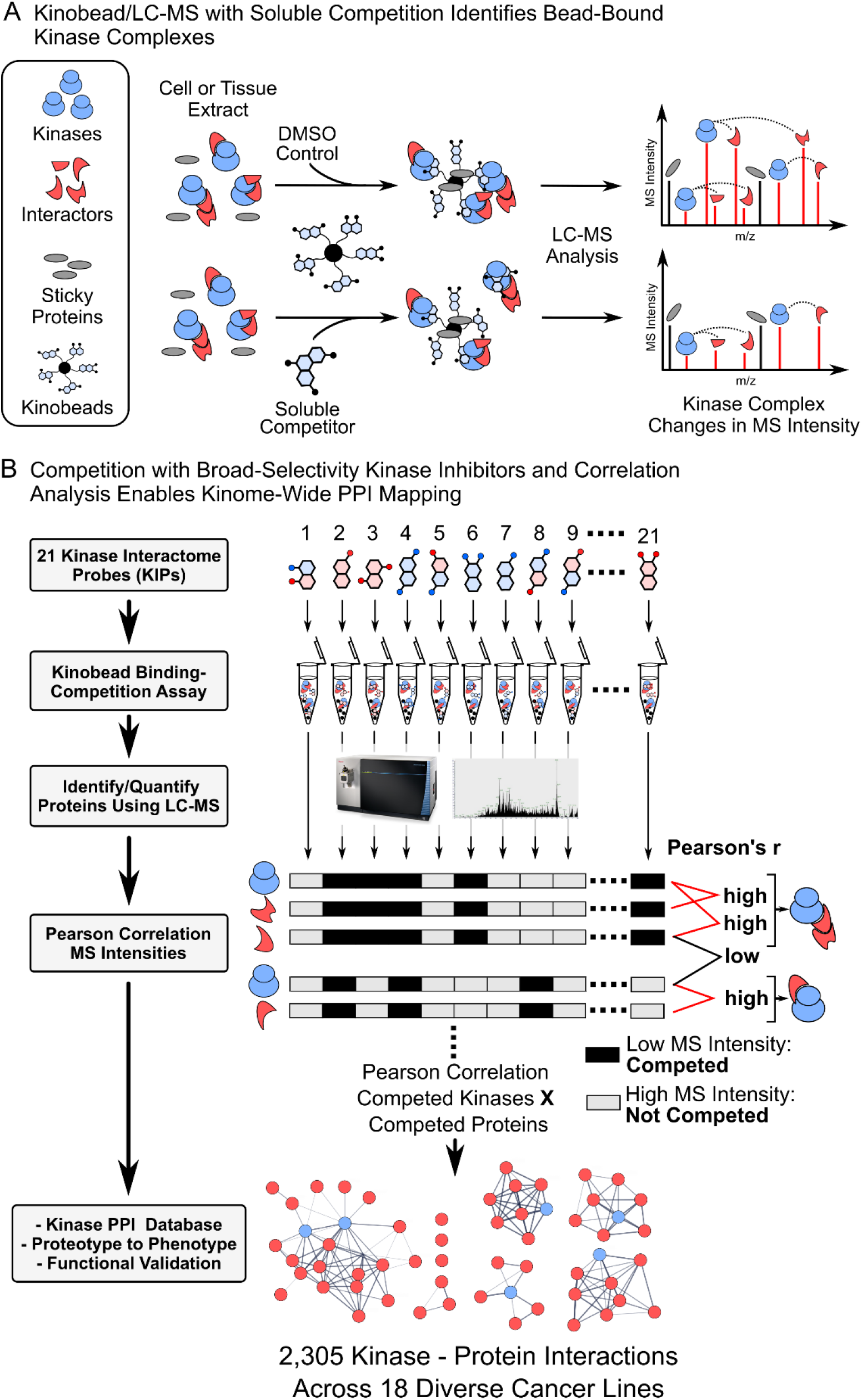
Overview of our kinobead competition and correlation analysis (Ki-CCA), a chemoproteomics approach for multiplexed, kinome-wide PPI profiling. **A)** Kinobead/LC-MS with kinase inhibitor (KI) soluble competition identifies bead-bound kinase complexes. **B)** Workflow of our Ki-CCA analysis using 21 kinase interactome probes (KIPs) that broadly compete kinobead-bound kinase complexes.

To find an optimal set of KIPs for our Ki-CCA, we mined kinobead profiling data of kinase-targeted drugs ^16^. We extracted the kinase MS intensity ratios from KI competed and DMSO control samples at the highest tested KI concentration (30 µM) and collated the data into a single matrix containing the ratios for 241 kinases competed with 220 KIs (Table S1). We then applied pairwise Pearson correlation of MS intensity ratios for all KIs and applied unsupervised hierarchical clustering to the resulting matrix of r-values, thereby identifying 12 clusters of KIs with similar binding profiles across the human kinome (Figure 2A and Table S1). To obtain a set of KIPs with complementary kinome binding affinities, we chose one to five KIs from clusters 1-5, 8-10, and 12 that contained the most broadly selective inhibitors (Figure 2A and Table S1). To validate that our set of 21 KIPs competes most kinases quantified by kinobead/LC-MS ^19,21^, we profiled their kinome selectivity using label-free quantification (LFQ) at a single high concentration in unstimulated HeLa cell lysate (10-50 µM, see ‘*Materials and Methods*’). This analysis quantified 224 kinases and showed that 199 of them (90%) were efficiently competed by at least one of our KIPs (log2 LFQ-MS Intensity ratio >0.75, two sample t-test p < 0.1, n = 2, Figure S1A and Table S2). We found that most kinases bind to 5-13 KIPs and that our probe panel uniformly competes kinases from all sub-families of the kinome, indicating that our probes are well suited to unbiasedly compete most kinobead-bound kinase complexes for identification by Ki-CCA (Figure S1A). Kinobead/LC-MS profiling of our 21 KIPs also revealed that highly homologous kinases, for instance AMPK1 and 2 (PRKAA1 and 2), or MST1 and 2 (STK3 and 4), show very similar binding profiles to our probes. Consequently, our Ki-CCA may not be able to distinguish the interactomes of such kinases. To overcome this problem, we combined kinases with similar KIP binding profiles (Pearson’s r ≥ 0.9) into 294 kinase groups that consist of one to four members, e.g., the four members of the FGFR family of receptor tyrosine kinases (RTKs) form the FGFR1-4 group (Table S2, and ‘*Materials and Methods*’). Accordingly, our Ki-CCA reports interactions between kinase groups and non-kinase proteins.

**Figure 2.**
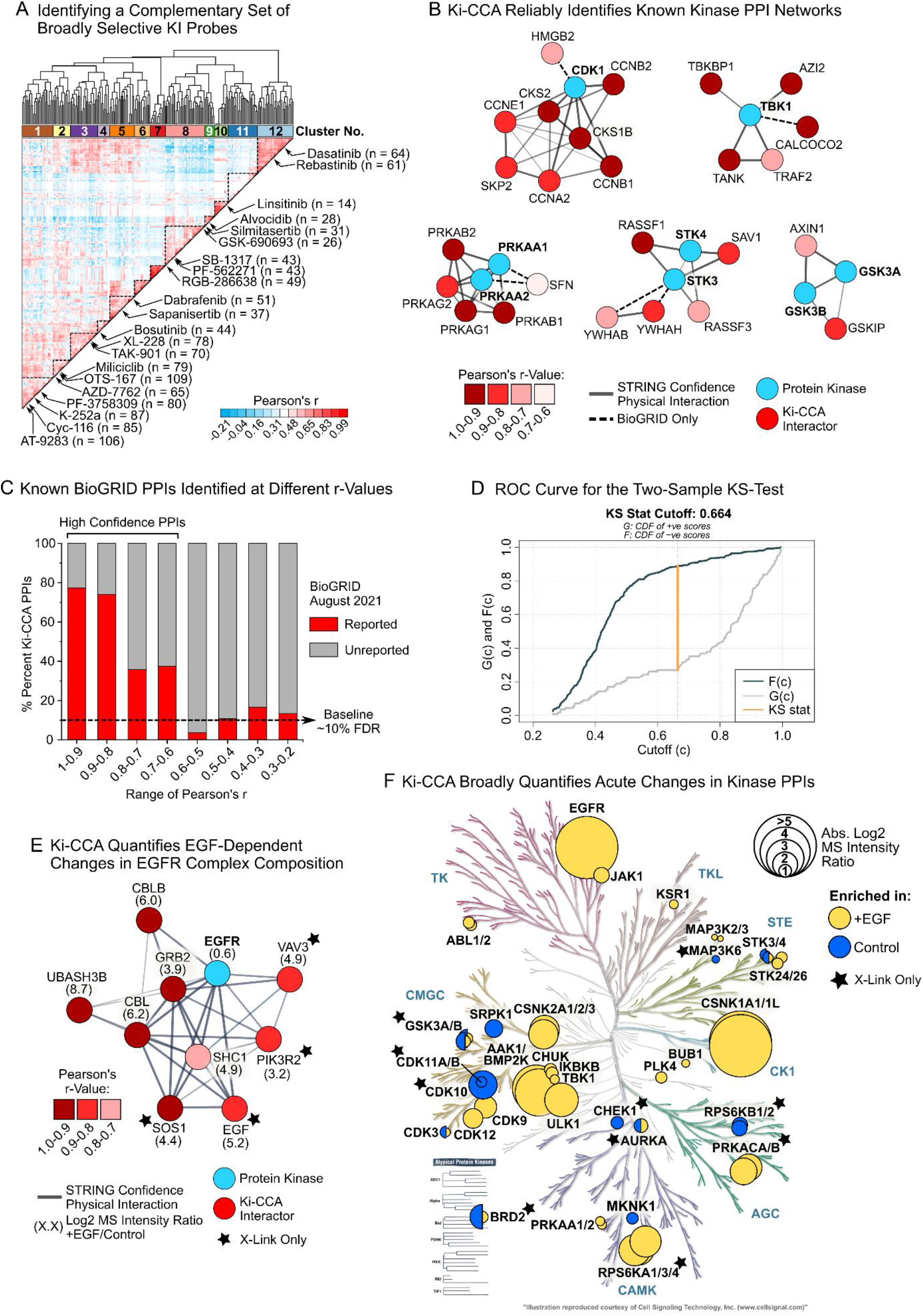
Selecting a panel of kinase interactome probes (KIPs) for Ki-CCA and proof-of-concept in HeLa cell lysate. **A)** Identifying broadly selective KIs with orthogonal kinome affinity to serve as kinase interactome probes (KIPs). **B)** Ki-CCA identified well characterized, stable kinase PPIs with high positive Pearson’s r-values. **C)** Percentage of Ki-CCA-predicted PPIs previously reported in the BioGRID and BioPlex 3.0 interaction databases, and the kinome interactomics study by Buljan et al. as a function of Pearson’s r-values (0.1-unit brackets). **D)** Example ROC plot of the two-sample KS-test for the unstimulated HeLa cell model (see also Figure S2). **E)** Ki-CCA quantifies changes in kinase PPI abundance in response to acute growth hormone treatment (HeLa cells + 15 min 50 ng/mL EGF). Shown is a ten-member activated EGFR complex. **F)** Significant kinome-wide changes in PPI abundance following acute EGF treatment in HeLa cells resolved by kinase. The kinase PPI with the highest absolute log2 MS intensity ratio was used for plotting (two sample t-test p < 0.1, n = 2 per condition).

### Ki-CCA identifies known as well as previously unreported kinase PPIs

To validate our Ki-CCA approach, we first tested if we could accurately identify known kinase PPIs. We and others have previously shown that kinobeads enrich multiple well-characterized, stable kinase complexes such as cyclin-dependent kinases (CDKs) associated with their cognate cyclins and other cell cycle proteins, and glycogen synthase kinase 3 (GSK3A and B) associated with AXIN1 ^16,17,19,22,23^. Expecting that Ki-CCA identifies such stable kinase PPIs, we subjected unstimulated HeLa cells to our workflow (Figure 1B and S1B). In addition to 199 competed kinases, we identified 573 competed non-kinase proteins that likely interact with these kinases. We correlated the MS intensity values of all competed kinases and non-kinase proteins, grouped kinases with similar KIP profiles (r > 0.9), and called the kinase group showing the highest r-value for each non-kinase protein its most likely interactor. Matching the resulting list of PPIs with the BioGRID protein interaction database (v.4.4.200, July 25 2021, primary and secondary kinase interactions)^24^, the BioPlex 3.0 dataset ^11^, and the kinome-centric AP-MS dataset from Buljan et al. ^10^, showed that 144 of 573 predicted interactions were previously known (Table S3). Among known PPIs we found, e.g., a CDK1 network comprising its regulatory subunits CKS1B and CKS2, and the cyclins A1, B1, B2 and E1. We also identified the CDK activating complex (CAK, CDK7-MNAT1-CCNH) and its interactions with components of the general transcription and DNA repair factor IIH (TFIIH) core complex (ERCC3 and 5, GTF2H1 and 4). Additionally, we identified PPIs between the NF-kB effector kinase TBK1, the signaling adapters TBKBP1, TANK, and AZI2 and the E3 ligaseTRAF2, as well as known interactions of 34 other kinase groups (Figure 2B and Table S3). These results confirm that Ki-CCA accurately and broadly identifies kinase PPIs.

Analyzing the distribution of Pearson’s r values for known interactions confirmed that Ki-CCA identifies PPIs with high positive values (median r of 0.83), and we observed a 3.5-fold increase in true positive identifications at r > 0.6 compared to the ∼10% baseline FDR (Figure 2C). Applying a two-sample Kolmogorov-Smirnov (KS) test to our dataset, comparing the r-value distributions of known and unreported PPIs, we found that at r-values ≥ 0.66 we preferentially sample true-positive PPIs (Figure 2D). Consequently, we defined a Ki-CCA r-value of ≥ 0.6 to identify high confidence kinase PPIs. Strikingly, applying this selection rule to our HeLa cell dataset, we identified high confidence interactions between 60 proteins and 21 kinase groups that have not been previously reported (Table S3). Among them we find, for e.g., interactions of CDK12 and CK2 (CSNK2A1-3) with the E3 ligases Arcadia (RNF111) and ARKadia-like protein 1 (ARKL1), respectively, linking these kinases to the ubiquitin-proteasome pathway. Collectively, our results demonstrate that Ki-CCA quantifies known kinase PPIs and identifies previously unreported interactions.

### Ki-CCA quantifies kinase PPI changes in response to acute signaling cues

Encouraged by our results, we investigated if Ki-CCA can quantify dynamic changes in kinase PPIs caused by activation of a specific signaling event, for e.g., growth hormone-mediated receptor activation. We stimulated HeLa cells with 50 ng/mL EGF for 15 min and subjected cell lysates to our Ki-CCA. We quantified 194 competed kinases and 540 competing non-kinase proteins, including 152 non-kinases that showed high confidence interactions with 44 kinase groups (r ≥ 0.6). Comparing protein MS intensity changes between unstimulated and EGF-stimulated HeLa cells revealed 277 non-kinase proteins with significantly altered abundance (two-sample t-test, p < 0.1, n = 2). These proteins may be part of kinase complexes whose composition is sensitive to EGF stimulation, and indeed, our Ki-CCA showed that 29 of these proteins interacted with 23 kinase groups (r ≥ 0.6). Among these PPIs we identified interactions between the activated EGFR and the signaling adapters GRB2 and SHC1, as well as the E3 ubiquitin ligases CBL, CBLB, and the tyrosine phosphatase UBASH3B (Figure 2E, Table S3). Furthermore, we identified multiple PPIs involving known downstream effectors of EGFR signaling, including members of the mitogen-activated protein kinase (MAPK) cascade such as MAP3K2/3 and KSR1, and the cell cycle kinases CDK9 and 12 (Figure 2F). These results suggests that Ki-CCA can map acute changes in kinase PPI networks, but the overall number of EGF-sensitive PPIs as well as their log2 MS intensity ratios appeared low given that EGFR signaling strongly affects at least 120 other kinases associated with numerous cellular pathways ^19,25^. Hypothesizing that Ki-CCA, like other AP-MS-based methods, disfavors the identification of transient and weak protein interactions, we repeated our analysis in EGF-stimulated and unstimulated HeLa cell lysates, this time applying formaldehyde-mediated protein crosslinking to stabilize kinase signaling complexes (see ‘*Materials and Methods*’). Indeed, Ki-CCA with protein crosslinking identified EGF-sensitive PPIs for 10 additional kinase groups, including interactions between the EGFR, the GTP-activating proteins SOS1 and VAV3, and the phosphatidylinositol 3-kinase regulatory subunit PIK3R2, as well as PPIs of known EGF signaling effectors such as the ribosomal S6 kinases RPS6KA1, 3 and B1, the cAMP-dependent protein kinase PKA, and the canonical WNT-pathway regulator GSK3B (Figure 2E, 2F and Table S3). We also found that crosslinking increased the number of identified EGF-sensitive PPIs per kinase by two-fold, i.e., an average of 3 compared to 1.5 interactors, and that the average absolute log2 ratios of interactors between stimulated and unstimulated cells increased from 1.4 to 2.0 (Table S3). Collectively, this shows that protein crosslinking stabilizes signaling complexes and that Ki-CCA with protein crosslinking can be a powerful approach for profiling dynamic changes of kinase PPIs in response to acute signaling cues.

### Ki-CCA Profiling of Diverse Cancer Cell Lines Reveals Kinase PPI Network Plasticity

Next, we investigated if our Ki-CCA can be used to identify kinase PPI networks that underlie the progression of different cancers. We analyzed 18 diverse cell lines representing distinct types of cancers, including carcinomas (10 HCC lines), adenocarcinoma (HeLa), blastomas, i.e., two neuroblastoma lines (SK-N-SH and SH-SY5Y) and one glioblastoma line (A172), myeloid malignancies such as CML (K562), MCL (JeKo1), and T-cell leukemia (Jurkat), and one osteosarcoma line (U2-OS, Table S2 and S3). Notably, many of these lines exist in distinct phenotypic states, for instance, at least three lines show mRNA expression signatures that are characteristic for epithelial-like cells, and seven lines show mesenchymal-like characteristics. This gives us the opportunity to study how cellular plasticity programs shape kinase PPI networks (Figure S1C, Table S2). Collectively, Ki-CCA profiling of these lines quantified 382 kinases (73% of the kinome) of which 357 were competed using our 21 KIPs. Concurrently, we quantified 6660 non-kinase proteins of which 4136 were competed. Our Ki-CCA correlation analysis predicted 10,791 PPIs between these proteins and 294 kinase groups, 1783 of these interactions having previously been reported in the BioGRID and BioPlex 3.0 databases ^11,24^, and a kinome-centric interactomics study by Buljan et al. ^10^. Analyzing the r-value distribution of known PPIs confirmed our observation in HeLa cells, i.e., that a ∼2-fold leap in true positive PPI identifications occurs at r-values ≥ 0.6 (Figure S3). This analysis also revealed a marked increase in true positive identifications at r-values of 0.5 to 0.6, leading us to define Ki-CCA-predicted PPIs in this r-value window as intermediate confidence interactions. Subjecting Ki-CCA interactions from the 18 cell lines to two-sample KS-tests showed that most true-positive PPIs were identified at a median r-value of ≥0.597, validating that a r ≥ 0.6 cut off reliably identified kinase PPIs. Applying these selection rules, we mapped 1054 high confidence 1051 intermediate confidence kinase PPIs (Table S3). Considering high and intermediate confidence interactions, each kinase group interacted on average with nine proteins, with numbers ranging from one interactor, e.g., for the nick-related protein kinase NRK, to up to 103 for casein kinase 2 (CK2 or CSNK2A1, 2, and 3, Table S3). This indicates that certain kinases preferentially form large, stable PPI networks that can be quantified with Ki-CCA.

Strikingly, we found that 691 of high confidence kinase PPIs (60%) have not been reported in our reference datasets, demonstrating that Ki-CCA is a powerful approach for discovering novel kinase interactions (Table S3). Furthermore, when we analyzed the recurrence of interactions across the 18-cell line panel, we found that 686 of high confidence PPIs (61%) were identified in only one cell line. This suggests that the composition of kinase complexes is highly context dependent, and that our Ki-CCA dataset can illuminate the pathobiology of individual cancer cell lines and other models that reflect the heterogeneity of cancer types and patients’ tumors. Indeed, analyzing high and intermediate confidence interactions that occur only in one type of cancer line, we identified PPI networks of kinases that are well-known to drive cancer progression or control important physiological functions. For instance, in the Philadelphia Chromosome positive (Ph^+^) CML line K562, Ki-CCA identified a 11-member PPI network centered on the BCR-ABL1 fusion protein, that included, e.g., the mitogenic signaling adapters SHC1, GRB2, the E3 ligase CBL, as well as the previously unreported interactor SCAI that regulates transcription and suppresses tumor cell migration (Figure 3A) ^26^. BCR-ABL1 is preferentially localized to the cytoplasm, suggesting that the fusion protein can sequester SCAI to block its tumor suppressor function in the nucleus ^27^. In EGFR-mutant A172 glioblastoma cells, Ki-CCA identified an activated six-member EGFR network composed the known interactors GRB2, SHC1, ERRFI1, and the sphingolipid transporter spinster 1 homolog (SPNS1), validating that the EGFR drives A172 cell proliferation and identifying a previously unreported EGFR interactor (Figure 3A). In the widely used Jurkat T-cell model, Ki-CCA identified several complexes of non-receptor tyrosine kinases that control T-cell receptor signaling and T-cell development. For example, we quantified a six-member network of LCK containing the known interactors MHC class II coreceptor CD4, the Src-family kinase (SFK) activating protein UNC119 as well as a previously unreported interaction with the myristoyl-binding protein and cargo adapter UNC119B. This suggests that UNC119B is an additional factor besides UNC119 directing LCK to the immune synapse ^28^. We also identified a seven-member CSK network that, besides its known interactor PAG1, also contained the putative Myc transcriptional activator PURA ^29^, the nuclear receptor corepressor 2 (NCOR2) that can control hematopoietic stem cell pools ^30^, and the E3 ubiquitin ligases UBASH3A and B. This gives novel insights into how CSK may regulate T-cell receptor recycling and T-cell development (Figure 3A) ^31^. Concurrently, we found that in non-hematopoietic cells CSK preferentially interacts with paxillin (PXN) and tensin 1 and 3 (TNS1 and 3), highlighting its function in focal adhesion signaling (Table S3). Together, our results demonstrate that Ki-CCA reliably quantifies kinase PPI networks that drive cancer progression, and important physiological processes, and identifies unreported PPIs that broaden our knowledge of these kinases.

**Figure 3.**
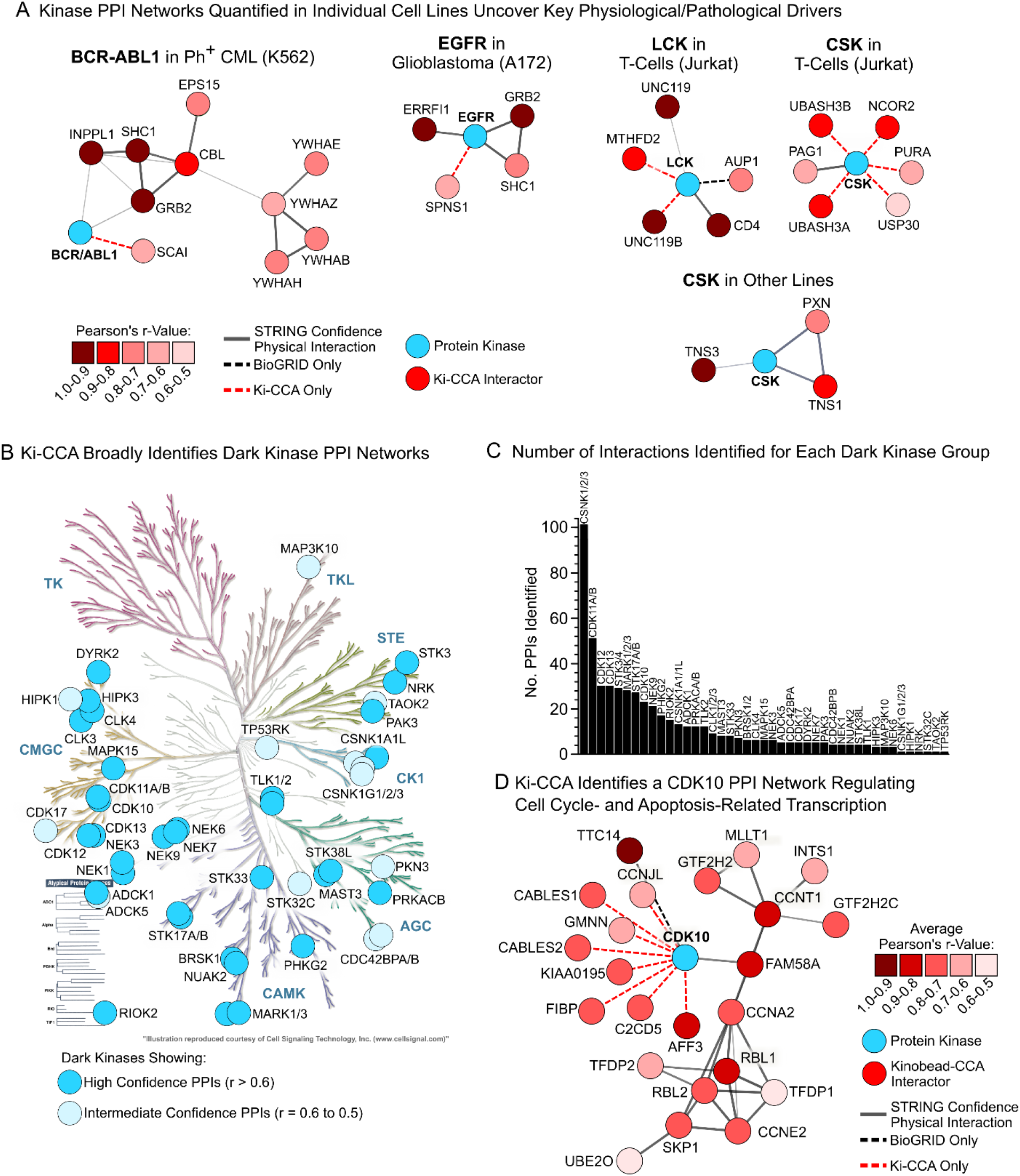
Ki-CCA identifies kinase PPI networks that control cancer progression and T-cell function, including dark kinase interactions. **A)** Example Ki-CCA-predicted kinase complexes that control important cellular phenotypes in specific cancer types. **D)** Dark kinases for which Ki-CCA identified PPIs overlayed with the human kinome dendrogram. **E)** Number of interactions for each of the dark kinase groups for which PPIs were identified. **F)** Ki-CCA PPI network for the dark kinase CDK10.

### Ki-CCA identifies “dark” kinase PPIs, shedding light on unknown biological functions

Despite their importance in physiology and disease, approximately 1/3 of all protein kinases are considered “dark” because their biological functions are virtually unknown ^32^. This is in part due to a lack of immunoprecipitation-grade antibodies that could be used to map dark kinase interaction networks. To learn if our Ki-CCA identified PPIs of dark kinases that may reveal their biological function, we aligned our PPI dataset with the list of kinases designated as ‘understudied’ by the NIH Illuminating the Druggable Genome (IDG) initiative (Table S3) ^33^. This revealed that our interaction database contains 536 high and intermediate confidence interactions between 43 dark kinase groups and 392 non-kinase proteins, covering 32% of the understudied kinome (Figure 3B). On average we find 12 interactors for each dark kinase group, and like their well-studied counterparts certain kinases form very large stable interaction networks, e.g., the CDK11A/B group (51 PPIs), followed by the CDK12, CDK13, STK3 and 4, and MARK1-3 group (25 to 30 PPIs) (Figure 3C). Remarkably, we identified a 24-membered CDK10 network, of which only two members, its cognate cyclin CCNQ (FAM58A) and the tetratricopeptide repeat protein 14 (TTC14), were previously reported, indicating that Ki-CCA greatly expands the knowledge of CDK10’s biological function (Figure 3D). The STAR syndrome-deficient kinase CDK10 ^34^ can act as a tumor suppressor or promote tumor progression depending on the type of cancer ^35^. We found that CDK10 interacted with the retinoblastoma-like proteins 1 and 2 (RBL1/2), two key regulators of cell cycle entry and apoptosis, as well as the transcription factors Dp-1 and Dp-2 (TFDP1 and 2) that control the expression of cell cycle genes and suppress differentiation ^36^. Furthermore, CDK10 interacted with the DNA replication inhibitor geminin (GMNN) and CDK5 and ABL1 enzyme substrate 1 and 2 (CABLES1 and 2) that promote CDK2 inactivation and p53-mediated apoptosis, collectively suggesting that CDK10 balances negative regulation of cell cycle progression with apoptotic stimuli. This is supported by our finding that CDK10 and CCNQ were preferentially expressed in slow-cycling mesenchymal-like HCC cells, possibly protecting these cells from replication stress observed in fast-cycling epithelial-like HCC cells ^22,37^. Together, these results demonstrate that our Ki-CCA PPI dataset can be used to illuminate dark kinases.

### Ki-CCA data enables mining of kinobead/LC-MS profiling data, identifying kinase PPI networks enriched in tumor tissues

Kinase interactome data of human tumor specimens could serve an invaluable resource to identify dysregulated signaling pathways and drug targets. Unfortunately, such data are difficult to obtain because it is challenging to express bait proteins for AP-MS in harvested tissues. Furthermore, IP-grade antibodies are unavailable for many kinases, limiting the feasibility of unbiased kinome-wide interactomics studies. Previously, we analyzed patient-derived HCC tissue samples using kinobead/LC-MS to identify kinases that are enriched in tumors, quantifying hundreds of kinases, and thousands of non-kinase proteins ^22^. We hypothesized that our Ki-CCA data of 18 diverse cancer cell lines can be used as a reference database to assign these proteins to specific kinase interaction networks. Consequently, we compared non-kinase proteins quantified in four paired HCC and normal adjacent liver specimens with our Ki-CCA dataset and identified 482 high confidence PPIs between 132 kinase groups and 244 non-kinase proteins present in the four HCC patient’s tissue samples (Table S3) ^22^. Quantifying the abundance differences of these kinase-interacting proteins between tumors and paired NAL showed that on average 183 kinase group-protein interactions were significantly altered (t-test with Benjamini-Hochberg (BH) FDR < 0.05, n = 5 or 6, Figure 4A, S4A and Table S3). Particularly, tumor 4 was highly enriched in PPIs of cell cycle kinases that are often aberrantly activated in cancers, for instance, CDK1, 2 and 7, and AURKB (Figure 4A and Table S3), indicating aggressive tumor growth. Particularly, tumor 4 was highly enriched in PPIs of cell cycle kinases that are often aberrantly activated in cancers, for instance, CDK1, 2 and 7, and AURKB (Figure 4A and Table S3), indicating aggressive tumor growth. Furthermore, PPI networks of kinases that control cell survival pathways (TBK1 and NUAK1), migration (DDR1 and 2, TNK2) and the Hippo pathway (STK3/4) were enriched in tumor 4, reflecting the high therapy resistance and invasiveness of HCCs. In contrast, the T-cell-specific LCK-CD4 interaction was enriched in normal adjacent liver tissue of case 4, suggesting poor immune cell infiltration of this tumor. The largest interaction network identified across the four patients’ tissues counted 41 members and was centered on CK2 (Figure 4B and Table S3). This CK2 network was highly enriched in components of the eukaryotic initiation factor 3 (eIF3) complex, the nucleosome remodeling deacetylase (NuRD) complex, and the polycomb-group (PcG) complex, as well as other components that control chromatin remodeling (DEK), the cell cycle, and proliferation (ERH, HDGFRP2). Notably, we found that CK2 does not change in abundance between tumors and NAL tissues but preferentially engages multiple eIF3 complex components and the NuRD complex component RBBP4 in three of four patient’s tumors (Figure 4C), suggesting that CK2 regulates both translation and chromatin remodeling in tumor tissues to promote aberrant growth. Based on these data, we propose that the clinical-grade CK2 inhibitor silmitasertib (CX-4945), either alone or in combination with other drugs, may be a useful new therapy in HCC ^38^. Collectively, our analysis of human tissue samples demonstrates that our Ki-CCA dataset acts as a powerful resource to identify kinase signaling complexes in existing kinobead profiling data.

**Figure 4.**
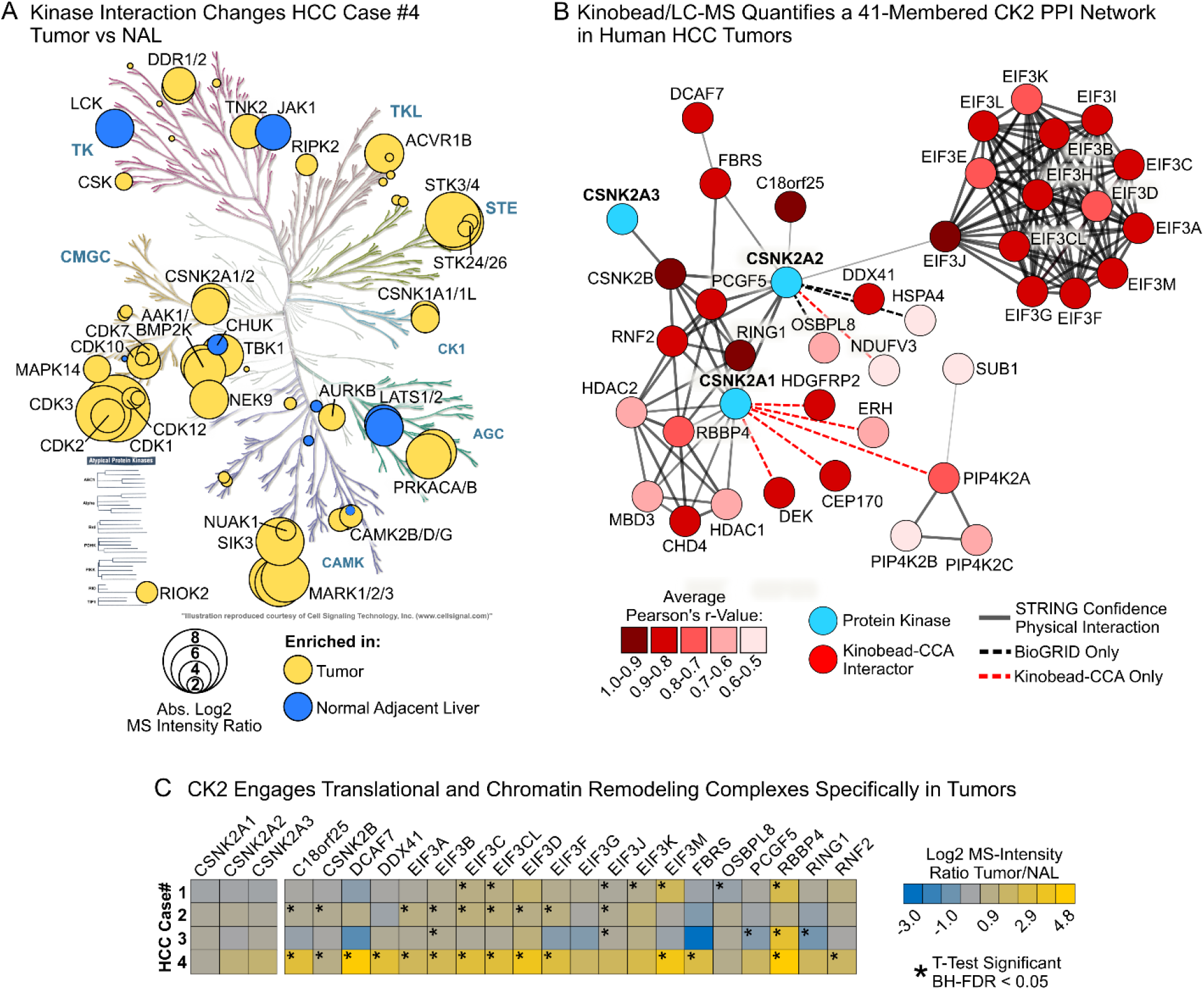
Using Ki-CCA data as a reference to identify kinase PPI networks in kinobead/LC-MS tissue profiling data. **A)** Overlay of the human kinome dendrogram with kinases whose interaction network composition significantly differed between human HCC tissue and normal adjacent liver (t-test BH-FDR <0.05). The kinase PPI with the highest absolute fold-change was used for plotting ^22^. **B)** Comparing our Ki-CCA database with kinobead/LC-MS profiling data identified a 41-member CK2 interaction network in human HCC tissue samples. **C)** Abundance differences of CK2 PPI network components between HCC patient’s tumor tissue and normal adjacent liver.

### AXL-dependent epithelial-mesenchymal transition globally rewires kinase PPI networks

Cancer cell EMT critically contributes to metastasis and therapy resistance in HCC and other types of cancers ^39,40^. We previously showed that the receptor tyrosine kinase AXL globally rewires kinome expression and phosphorylation to promote HCC cell EMT ^22^. Our kinobead/LC-MS studies also showed that *AXL* RNAi knockdown causes significant abundance changes in 887 non-kinase proteins, many of them known kinase interactors. We hypothesized that these proteins are part of kinase complexes that contribute to EMT and may therefore present attractive novel drug targets. To identify the kinase interactors of these EMT-associated proteins, we applied Ki-CCA to our FOCUS cell *AXL* RNAi model and FOCUS WT cells (Table S3). Our analysis revealed 106 high confidence and 38 intermediate confidence PPIs between 61 kinase groups and 123 non-kinase proteins whose abundance was sensitive to *AXL* knockdown (Figure 5A and S4B). Among them, we identified several PPIs that act on well-known EMT signaling pathways, such as GSK3B-AXIN1 that promotes β-catenin degradation and WNT signaling (2.3-fold increase), TBK1-TBKBP1 that activates NF-kB (3.4-fold decrease), and STK3/4-SAV1 that decreases Hippo/YAP-TAZ activity (3.5-fold decrease)^41^. This suggests that AXL signaling promotes WNT and NF-kB signaling and counteracts Hippo/YAP-TAZ activity. Strikingly, 21 kinase PPI networks with previously unreported roles in EMT were highly enriched in mesenchymal FOCUS WT cells. For instance, an interaction of CSK and the SH2-domain containing protein tensin-3 (TNS3, 7.5-fold decreased) indicated that CSK is tethered to focal adhesion complexes specifically in mesenchymal cells. We found that HCK, another SFK, interacted with galectin-1 (LGALS1) that induces T-cell apoptosis and activates HRAS, exclusively in mesenchymal-like cells, suggesting that this kinase may play a role in immune evasion and mesenchymal-like cancer cell proliferation. Components of a 27-membered CK2 network were enriched either in epithelial-like cells or in mesenchymal-like cells, among the latter the tumor suppressor and arginine N-methyl transferase effector EPB41L3 ^42^ (Figure S4B and Tables S3), establishing a connection between CK2, protein N-methylation, and EMT. Notably, several components of an eleven-member network of the AAK1/BMP2K kinase group was highly enriched in mesenchymal cells, including the small GTPase effectors RALBP1, REPS1, and REPS2, that showed 2.4-, 17-, and 9.2-fold decreased abundance following *AXL* knockdown (Figure 5B). This suggests a prominent role of this AAK1/BMP2K network in HCC cell EMT. AAK1/BMPK components are also significantly enriched in ten mesenchymal-like HCC cell lines compared to seven epithelial-like lines and overexpressed in two of four HCC patient’s tumors that we previously profiled using kinobead/LC-MS (Figure 5C and 5D) ^22^. Collectively, our results demonstrate that Ki-CCA identifies and quantifies known as well as previously unreported kinase PPI networks that may be important for cancer cell phenotypic plasticity and therapy resistance. Our results also indicate that an AAK1/BMP2K PPI network of unknown biological function may contribute to HCC cell EMT *in vitro* and cancer progression *in vivo*.

**Figure 5.**
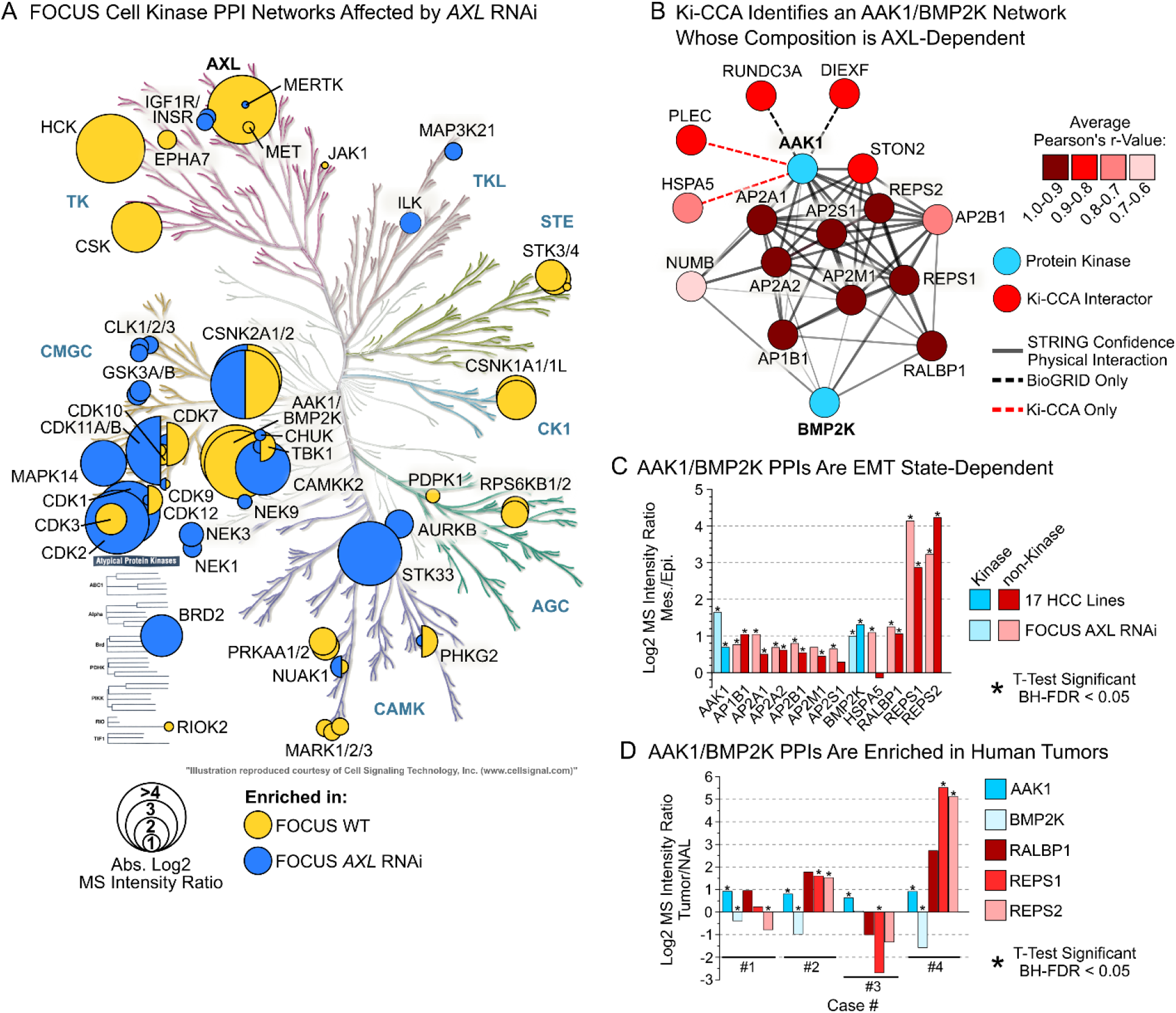
Ki-CCA profiling of AXL-dependent kinase PPI network rewiring identifies an AAK1/BMP2K network associated with EMT and tumor progression. **A)** Kinome dendrogram overlayed with FOCUS cell kinases whose PPIs significantly change in response to *AXL* RNAi knockdown (BH-FDR < 0.05, n = 5). The PPI with the highest fold-change was used for potting. **B)** Ki-CCA interaction network of high confidence interactors (r > 0.6) for the AAK1/BMP2K kinase group identified across the 18-cell line panel. **C)** Abundance differences of AAK1/BMP2K network components between FOCUS WT/*AXL* RNAi cells, and mesenchymal-like and epithelial-like cells of a 17 HCC line panel (kinobead/LC-MS data). **\*** t-test significant, BH-FDR < 0.05. **D)** Abundance differences of AAK1/BMP2K complex components between human HCC tumor tissue and normal adjacent liver (kinobead/LC-MS data). **\*** t-test significant, BH-FDR < 0.05.

### An AAK1-RALBP1-REPS1/2 complex promotes HCC cell EMT and therapy resistance

Having linked an AAK1/BMP2K interaction network to cancer cell EMT, we wished to study its potential as a drug target to overcome metastasis and therapy resistance. AAK1 plays key roles in clathrin-mediated endocytosis ^43^, whereas the function of BMP2K is less well understood.

More recent studies have shown that AAK1 is also involved in canonical Notch and Wnt/Fzd signaling that can contribute to cancer cell EMT ^44,45^. RALBP1 functions in Ral-mediated receptor endocytosis, serving as a GTPase activating protein (GAP) for CDC42/Rac ^46,47^. RALBP1 is also known to act as a drug efflux pump ^48^. Similar to RALBP1, REPS2 regulates growth hormone receptor endocytosis downstream of Ral ^49^, as well as cell migration and NF-κB pathway activation ^50,51^. In contrast, specific biological functions for REPS1 are not known. First, to clarify if either AAK1 or BMP2K is the central kinase of the network, we performed a kinobead/LC-MS soluble competition experiment in FOCUS cell lysate using 1 µM of the selective AAK1 inhibitor LP-935509 (Figure 6A and Table S4) ^52^. Our results show that RALBP1, REPS1 and 2, as well as four members of the adapter protein (AP) complex were competed at ratios similar to AAK1 (∼10-100-fold), whereas BMP2K was competed less than 4-fold, suggesting that the complex is centered on AAK1. To validate this finding, we performed Co-IP/MS experiments using antibodies targeting AAK1, RALBP1 and REPS1, and a GFP antibody as the control in FOCUS cell lysate (Figure 6B). These experiments showed that AAK1, RALBP1, and REPS1/2 co-precipitate with all three antibodies, validating that the network is centered on AAK1, and demonstrating that these proteins are part of the same AAK1 complex. Next, we disrupted the function of all four AAK1 complex components in four different mesenchymal-like and drug resistant HCC lines using shRNA-mediated gene knockdown (*‘Materials and Methods’*). Testing knockdown efficiency by qPCR, western blotting, and kinobead/LC-MS profiling showed that we achieved near-complete knockdown of RALBP1 and REPS1 and 2 (Figure S5A, B, and C). In contrast, AAK1 decreased in expression 2-4-fold, which is consistent with its critical function in endocytosis, an essential cellular process that likely prohibits complete removal of AAK1. Western blot analysis of five different EMT markers, i.e., AXL, CD44, CDH1 (E-cadherin), ZEB1, and SNAI1 (Snail), showed that at least two out of four RNAi lines experience drastic changes in EMT marker expression (Figure 6C and S6). Particularly, AAK1 complex RNAi in SNU387 cells nearly abolished AXL, CD44, and ZEB1 expression, and RNAi in FOCUS cells, especially REPS1 and 2 RNAi, greatly reduced CD44 and ZEB1 expression, and increased CDH1 expression. This suggests that AAK1 complex RNAi caused wide-spread EMT reversal in SNU387 and FOCUS cells.

**Figure 6.**
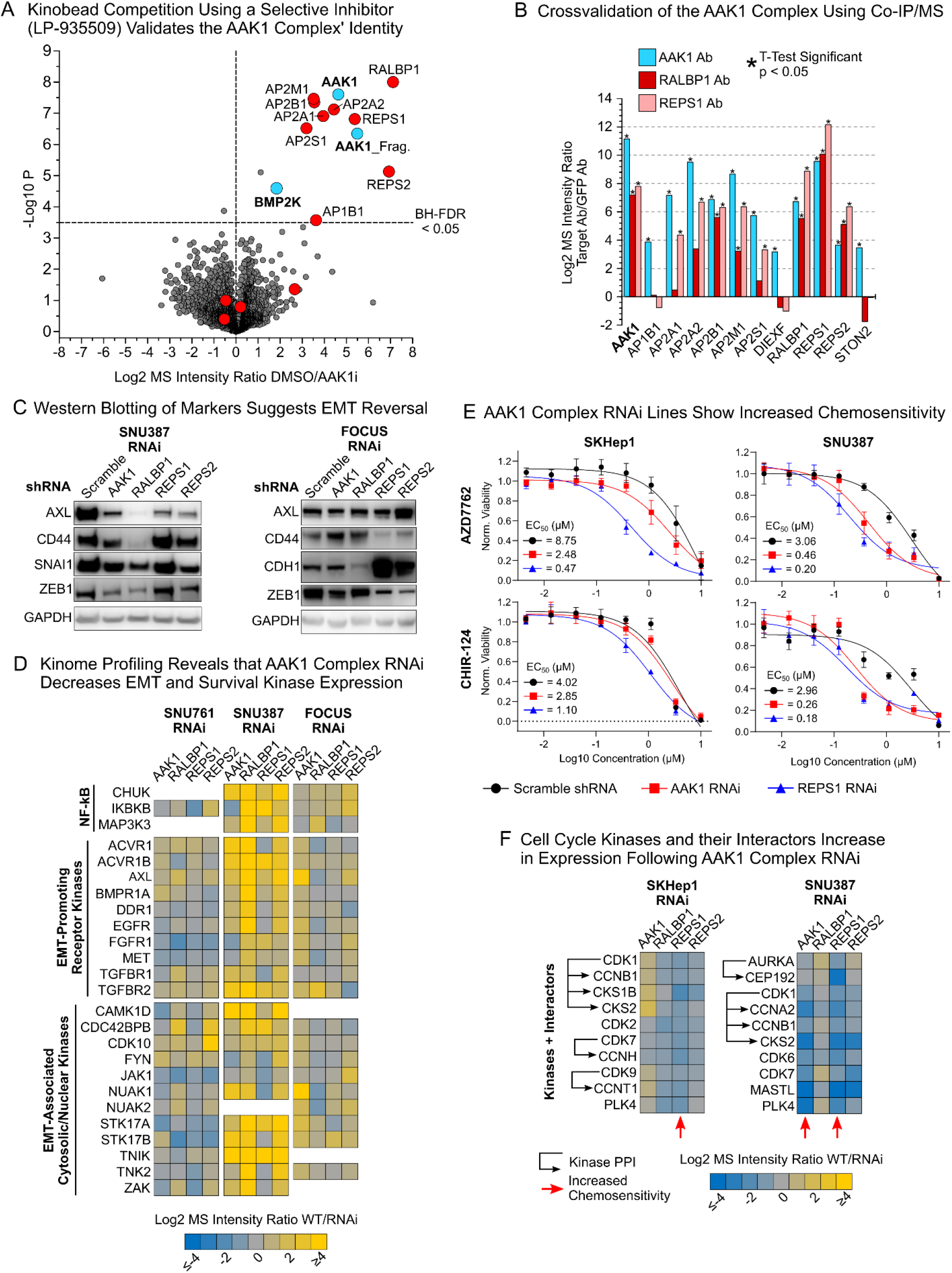
An AAK1-RALBP1-REPS1/2 complex promotes HCC cell EMT and therapy resistance. **A)** Kinobead/LC-MS soluble competition experiment using 1 µM of the selective AAK1 inhibitor LB-935509. Members of the AAK1/BMP2K PPI network are highlighted. **B)** Co-IP-MS experiments in FOCUS cells using antibodies targeting AAK1, RALBP1, and REPS1 validate the AAK1 complex’ identity. **C)** Western blot analysis of EMT markers in SNU387 and FOCUS RNAi lines. For analysis of the SKHep1 and SNU761 lines see Figure S6. **D)** Kinobead/LC-MS profiling data of AAK1 complex RNAi lines showing that NF-kB and EMT kinases consistently decrease in expression in three out of four cell line models. **E)** Drug screen results demonstrating that AAK1 complex RNAi lines are up to 18-fold more sensitive to the CHEK1 and 2 inhibitors AZD-7762 and CHIR-124. **F)** Kinobead/LC-MS profiling from SKHep1 and SNU387 lines showing that AAK1 complex RNAi activates cell cycle-related kinases.

To identify the mechanism of AAK1 complex RNAi-mediated EMT reversal, we mined our kinobead/LC-MS profiling data of RNAi lines for evidence of EMT pathway alterations (Figure 6D and Table S4). This revealed that AAK1 complex RNAi drastically decreases the expression of NF-kB effector kinases (IKBKG, MAP3K3, and CHUK) and numerous receptor tyrosine and serine/threonine kinases, including AXL, MET, EGFR, and TGFBR1 and 2 in out of four RNAi lines. This suggests that the AAK1 complex stabilizes membrane expression or recycling of receptor kinases that promote EMT pathway activation. Additionally, we find that many EMT-associated cytosolic and nuclear kinases decrease in expression in response to AAK1 complex RNAi, further supporting that knockdown causes EMT reversal (Figure 6D).

Next, to learn if RNAi sensitizes cells to drug treatment, we conducted a screen for cell viability with ten kinase-targeted drugs and doxorubicin, comparing responses of scramble shRNA lines with AAK1 complex RNAi lines (Figure 6E and TableS4). Remarkably, this drug screen showed that RNAi lines, particularly SNU387 and SKHep1, are up to 18-fold more sensitive to treatment with inhibitors targeting the DNA damage checkpoint kinases CHEK1 and 2 (AZD7762 and CHIR-124). To rationalize the increased efficacy of DNA damage response (DDR) kinase inhibitors, we analyzed kinome proteomics features that were upregulated in response to AAK1 complex RNAi in the cell lines with the greatest chemo-sensitization, i.e., SKHep1 and SNU387. We observed that these lines showed increased abundance of cell cycle-related kinases and their activated signaling complexes, notably, AURKA in complex with the activating scaffold CEP192, CDK1 complexes containing the cyclins CCNA2, E1, and B1, and the regulatory subunit CKS2, CDK6 and 7, PLK4, and CHEK1 (Figure 6F). These results indicate that RNAi lines, particularly AAK1 and REPS1 knockdown lines, contain more cycling cells that activate DDR signaling to enhance cell survival, and therefore are more sensitive to CHEK1/2 inhibition. ^53^. In contrast, SNU761 and FOCUS RNAi cells that show less chemo-sensitization also do not upregulate cell cycle-related kinases (Table S4). Furthermore, the SNU387 and SKHep1 REPS1 RNAi lines that show the greatest chemo sensitization exhibit the least alterations in EMT marker and kinase expression (Figure 6C and E, Table S4). These results demonstrate that knocking down components of the AAK1-RALBP1-REPS1/2 complex can cause downregulation of receptor and non-receptor kinases that promote EMT, thereby inducing EMT reversal. Concurrently, AAK1 and REPS1 knockdown can cause re-activation of the cell cycle and chemo sensitization, seemingly independent of EMT state. Collectively, our findings establish the AAK1 complex as promising drug target candidate to overcome cancer metastasis and therapy resistance.

## DISCUSSION

We introduced Ki-CCA, a chemoproteomics approach for highly multiplexed interactome mapping of the kinome. Ki-CCA allows the high-throughput profiling of various cell states and model systems using native cell and tissue lysates, entirely avoiding the use antibodies and the need to express genetically tagged bait proteins. We demonstrated that our Ki-CCA broadly quantifies kinase interactome changes in response to genetic manipulation of oncogenes, establishing Ki-CCA as a powerful approach for cancer research. Furthermore, we showed that Ki-CCA combined with protein crosslinking captures the rapid rearrangement of transient and low-affinity signaling complexes caused by acute stimuli, showcasing our approach’s utility for cell signaling research. Importantly, we demonstrated that our kinase PPI database can be used to identify kinase signaling complexes in kinobead/LC-MS kinobead profiling data derived from human tumor tissue samples, yielding novel insight into kinase PPI networks alterations in patient’s tumors and a roadmap to obtaining similar kinobead-based PPI networks in various disease models for clinical proteomics.

We demonstrated that Ki-CCA can help illuminate mechanisms of cancer therapy resistance and that it enables cancer drug target discovery. Our AAK1 complex example showcases that Ki-CCA can identify non-kinase target candidates such as REPS1 whose RNAi is better tolerated than AAK1 RNAi, strongly sensitizing mesenchymal-like cancer cells to targeted therapy. Therefore, inhibiting the AAK1-REPS1 interaction may be a promising strategy to minimize drug cytotoxicity and maximize cancer therapy responses. Ki-CCA combined with kinome profiling of RNAi lines showed that AAK1 and REPS1 knockdown reactivated the cell cycle. This reactivation was sufficient to sensitize HCC cell CHEK1/2 inhibitors, and that this effect was independent from changes in the cellular EMT state. Collectively, this indicates that the AAK1-REPS1 complex acts as a tumor suppressor, and that inhibition of tumor suppressors can be sufficient to sensitize mesenchymal-like cells to drugs ^54^. Consequently, complete reprogramming of mesenchymal-like cells to an epithelial-like state may not be required to achieve therapeutic benefit.

Finally, the limitations of our approach are the same as for any other AP-MS-based method, for instance, difficulties identifying weak and transient PPIs, and the lack of sub-cellular spatial resolution ^9^. Here, we already demonstrated that Ki-CCA with protein crosslinking can increase the coverage of weak and transient PPIs. Furthermore, we speculate that subcellular fractionation, e.g., into cytosolic, membrane, and nuclear fractions, for Ki-CCA could further resolve the localization of kinase signaling complexes and reduce sample complexity to increase the number of identifiable signaling and transcription factor complexes. Finally, the throughput of Ki-CCA can be further improved in future iterations of the approach by using smaller sets of KIPs and isobaric TMT labeling for analysis of entire interactomes in single LC-MS runs.

In summary, we presented a new approach for studying kinase interactome dynamics in virtually any model system. We collated our kinase interactome data into a database that is accessible through an interactive Shiny web application (https://quantbiology.org/kiCCA), serving as an important resource for cancer and cell signaling researchers.

## Supporting information

Identifying a complementary set of kinase interactome probes (KIPs).

Competed kinases in HeLa cells and EMT mRNA markers.

Summary kinase interactions 18 cell lines.

Kinobead-LC-MS profiling AAK1 complex and RNAi knockdown lines.

## ACKNOWLEDGMENTS

This work was supported by grants from the National Institutes of Health issued under the award numbers R01GM129090 (S-E.O.), R03TR003308 (M.G.), and R01GM086858 (D.J.M). This work used an EASY-nLC1200 UHPLC and Thermo Scientific Orbitrap Fusion Lumos Tribrid mass spectrometer purchased with funding from a National Institutes of Health SIG grant S10OD021502 (S-E.O.). The content is solely the responsibility of the authors and does not necessarily represent the official views of the National Institutes of Health.

## AUTHOR CONTRIBUTIONS

Conceptualization, M.G., S-E.O.; Methodology, M.G.; Investigation, M.G., A.L., T.S., H-T.L., and T.M.; Formal Analysis, M.G., S-E.O.; Writing – Original Draft, M.G., S-E.O.; Writing – Review and Editing, M.G., D.J.M, and S-E.O.; Funding Acquisition, S-E.O., D.J.M., and M.G.

## DATA AND CODE AVAILABILITY

MS .raw files, MaxQuant output files, and Ki-CCA correlation matrices generated by this study have been uploaded to the MassIVE repository of the University of San Diego under the acquisition number MSV000088067. A Shiny app for real time interrogation of the kinase interactomics data generated in this study can be accessed at https://quantbiology.org/kiCCA. This study did not generate new code.

## Supplementary Figures

**Figure S1.**
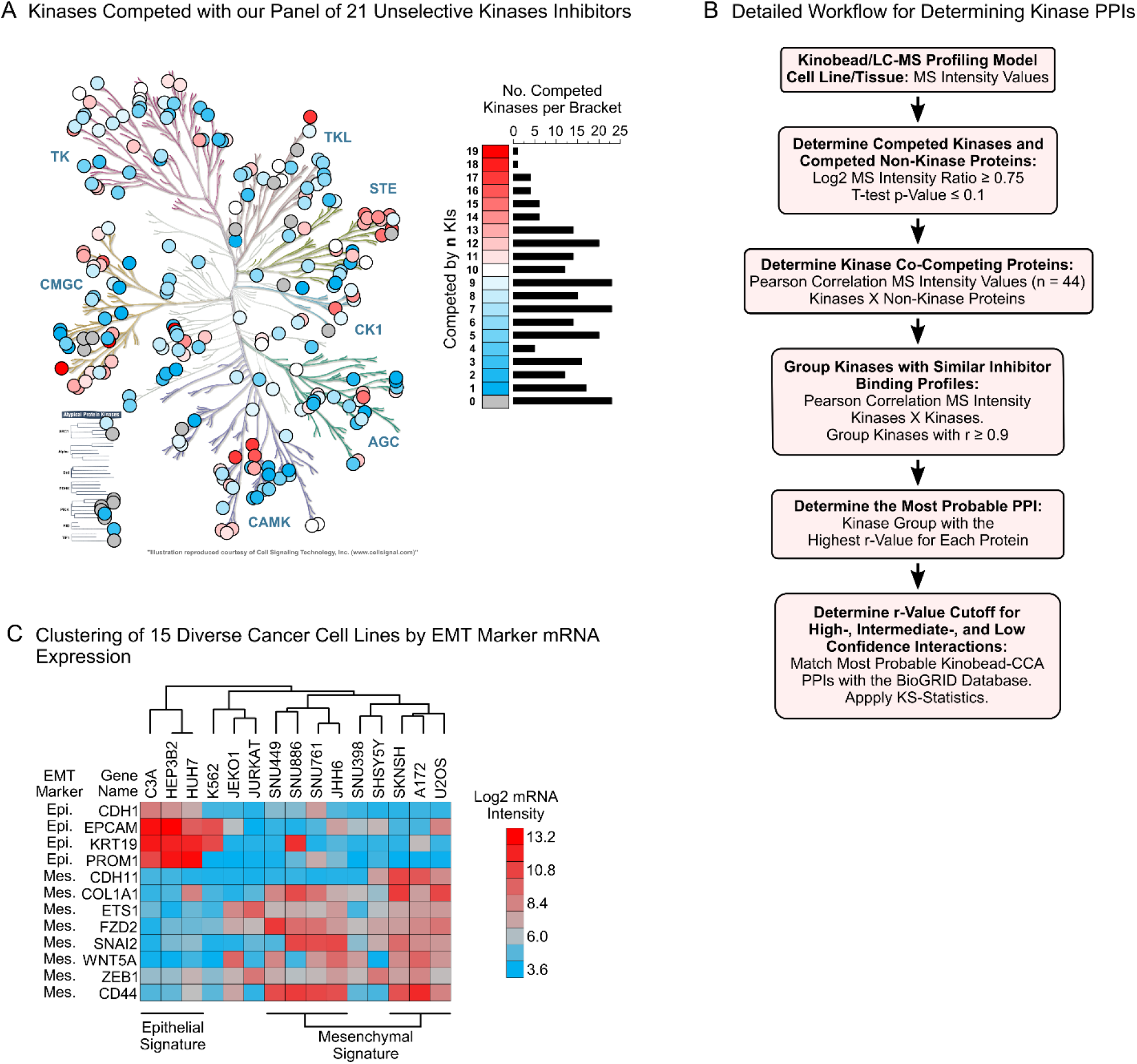
**A)** Kinases significantly competed in our kinobead/LC-MS soluble competition assay using our 21 KIP panel (HeLa lysate; log2 MS intensity ratio >0.75 and t-test p-value <0.1, n = 2). **B)** Detailed, step-by-step description of our Ki-CCA workflow to identify kinase PPIs. **C)** Clustering of 15 diverse cancer cell lines by EMT marker mRNA expression (n = 52). Shown are the 12 most characteristic markers for the epithelial and mesenchymal phenotype.

**Figure S2.**
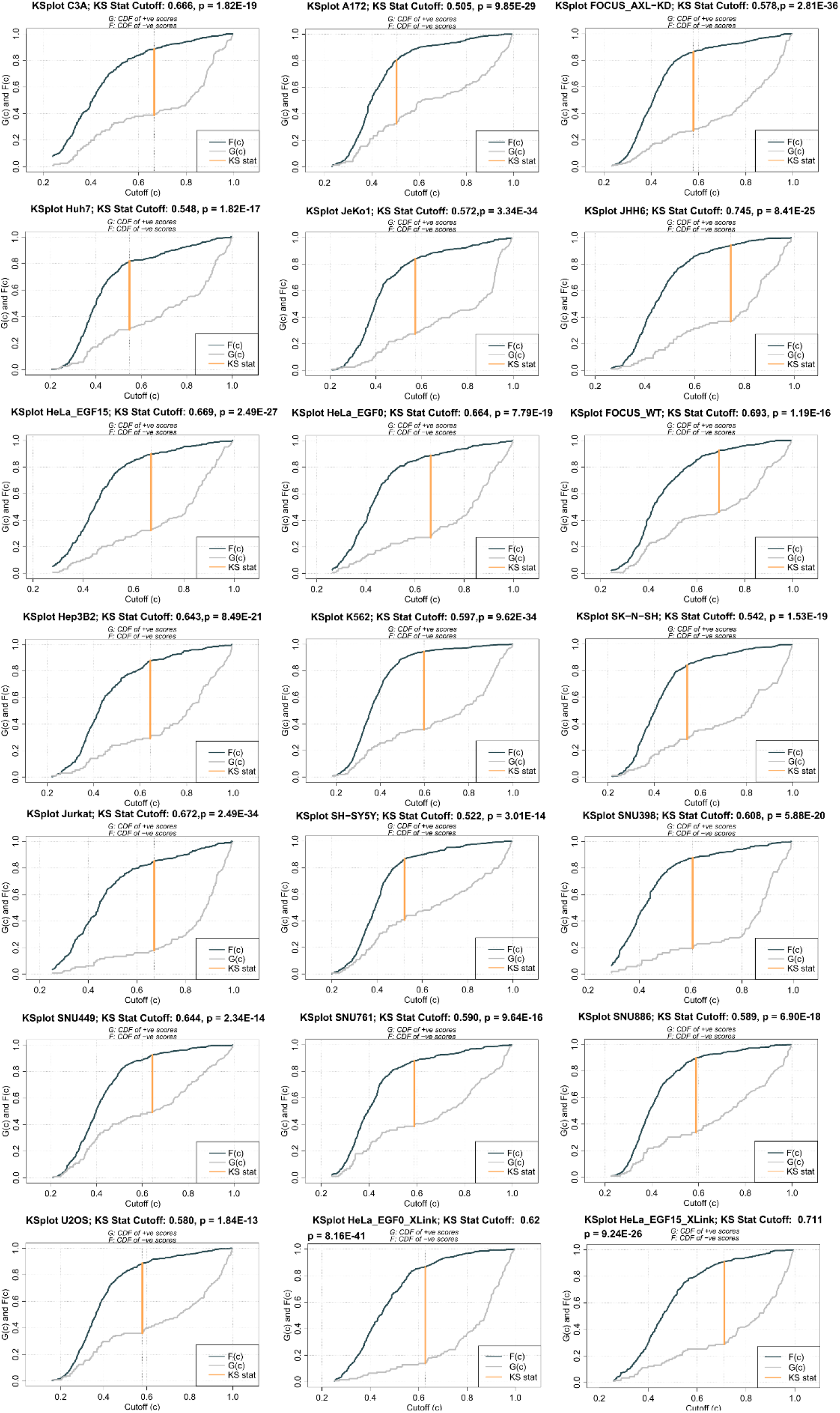
KS statistics for Ki-CCA profiling results of our 18 cell line models.

**Figure S3.**
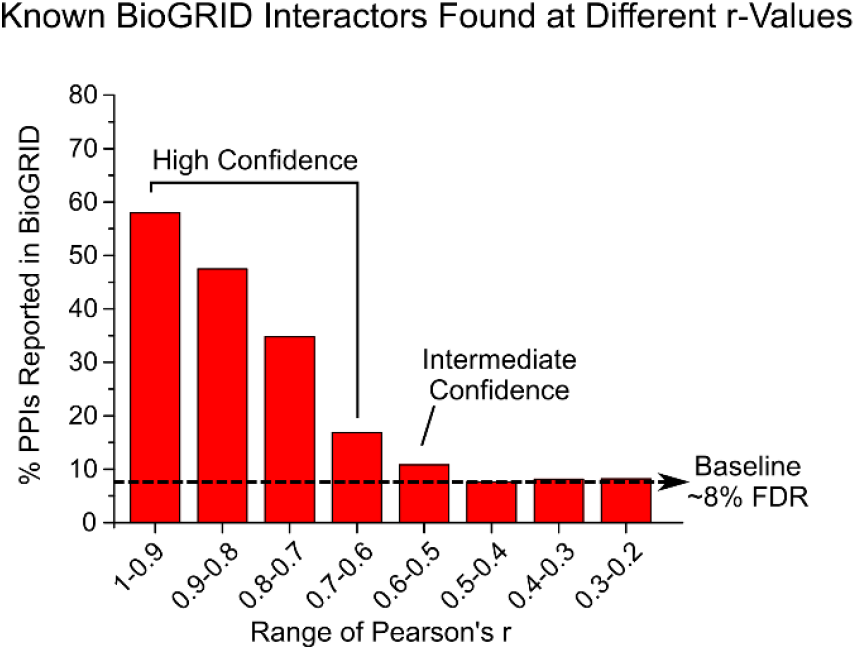
Percentage of Ki-CCA predicted PPIs previously reported in the BioGRID, BioPlex 3.0, and Buljan et al., 2020 interaction database as a function of Pearson’s r-values (0.1-unit brackets).

**Figure S4.**
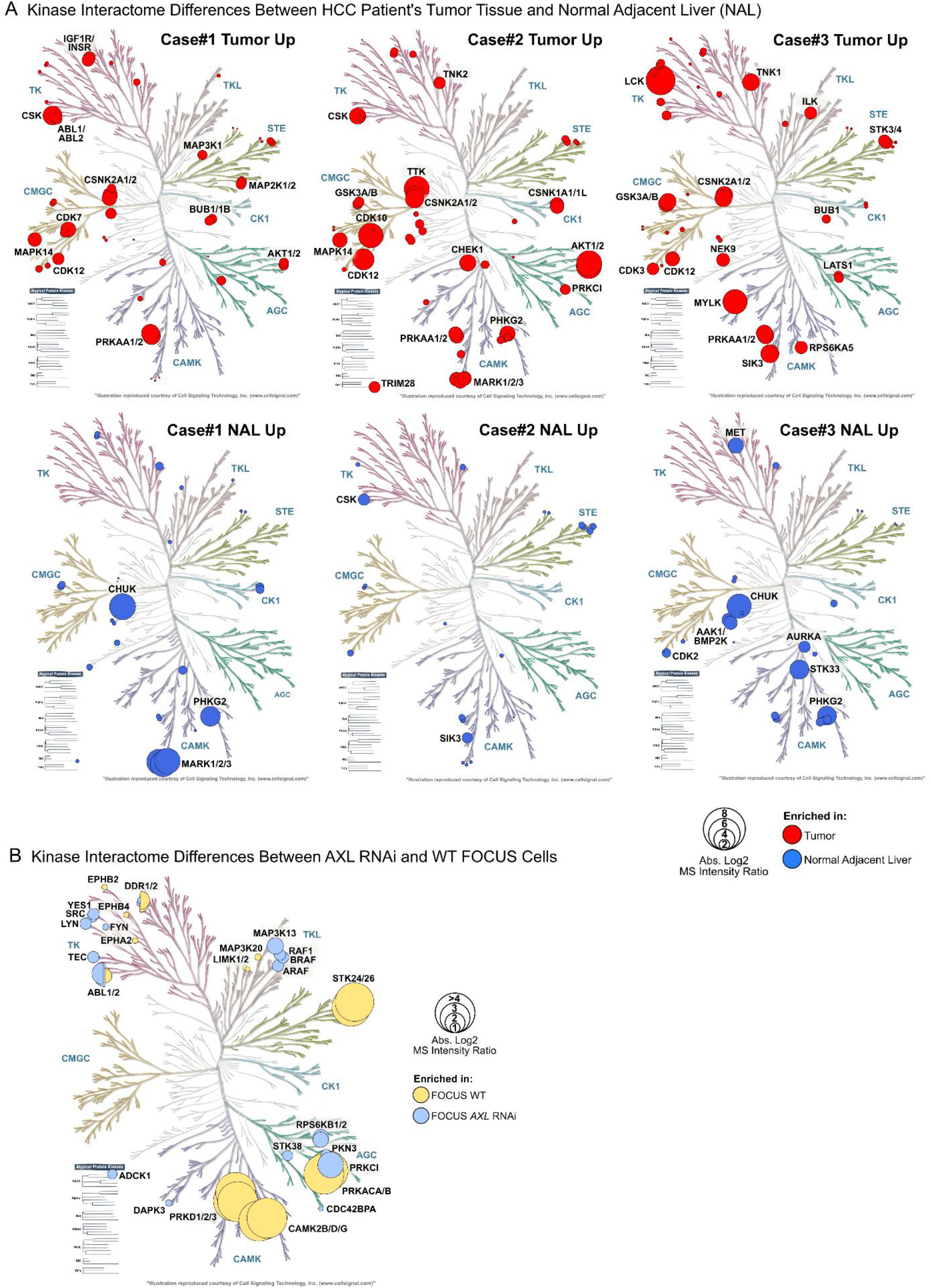
Kinase Interactome dynamics in human HCC patients’ tissues and a FOCUS *AXL* RNAi model of EMT. **A)** Overlay of the human kinome dendrogram with kinases whose complex composition significantly differs between human HCC tissue and normal adjacent liver (t-test BH-FDR <0.05). The kinase PPI with the highest absolute fold-change was used for plotting. **B)** Kinome dendrogram overlayed with FOCUS cell kinases whose PPIs significantly change in response to *AXL* RNAi knockdown (BH-FDR < 0.05, n = 5). The PPI with the highest log2 MS Intensity ratio was used for plotting. Only intermediate-confidence PPIs are shown (Ki-CCA r-value between 0.5 and 0.6).

**Figure S5.**
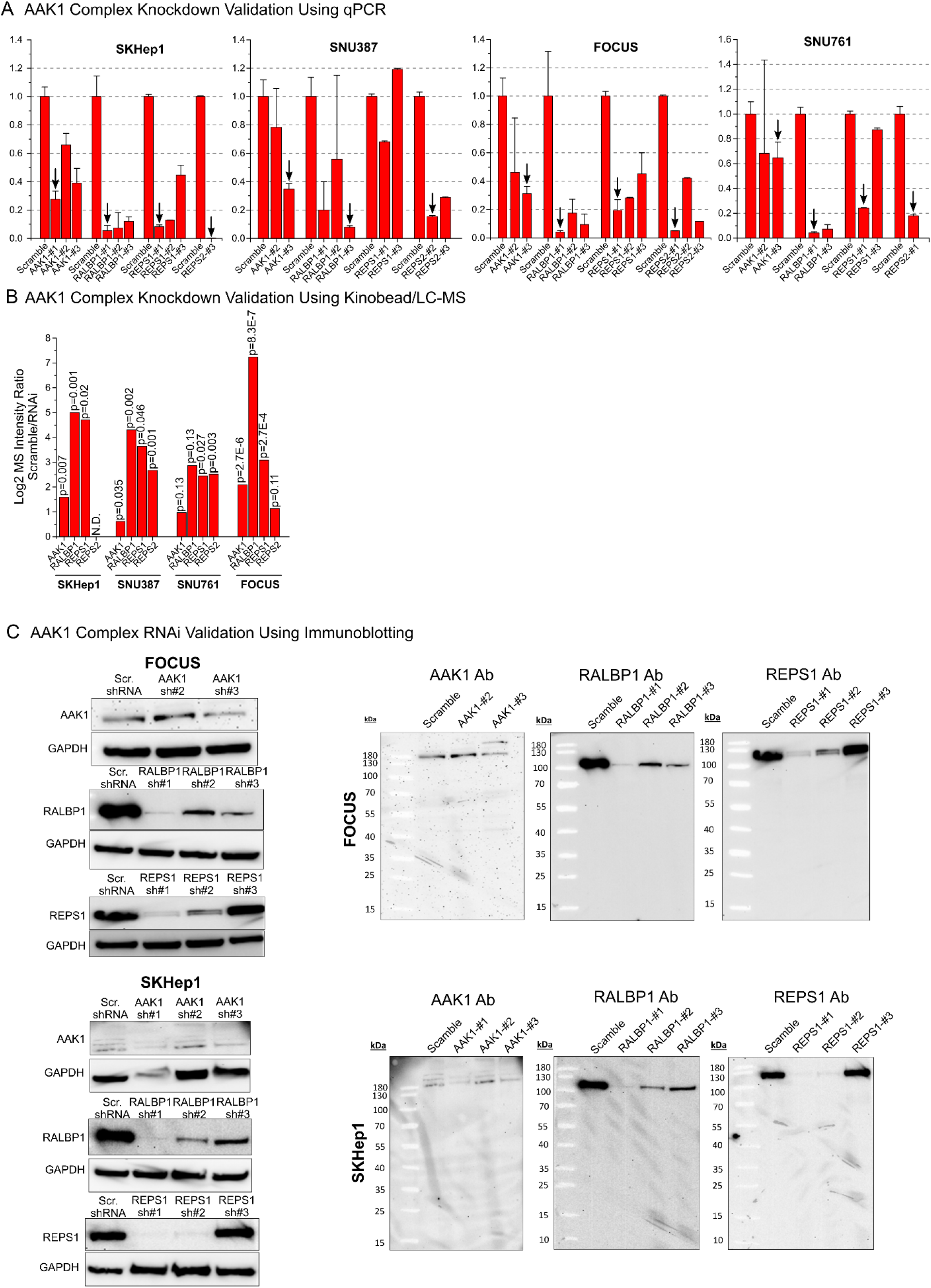
Validating AAK1 complex RNAi knockdown in FOCUS, SKHep1, SNU761, and SNU387. **A)** qPCR analysis of AAK1 complex RNAi lines, validating successful knockdown. **B)** Kinobead/LC-MS analysis of AAK1 complex RNAi lines, validating successful knockdown. **C)** Immunoblot analysis of AAK1 complex RNAi lines, validating successful knockdown. REPS2 blots not shown because the antibody used is likely not specific.

**Figure S6.**
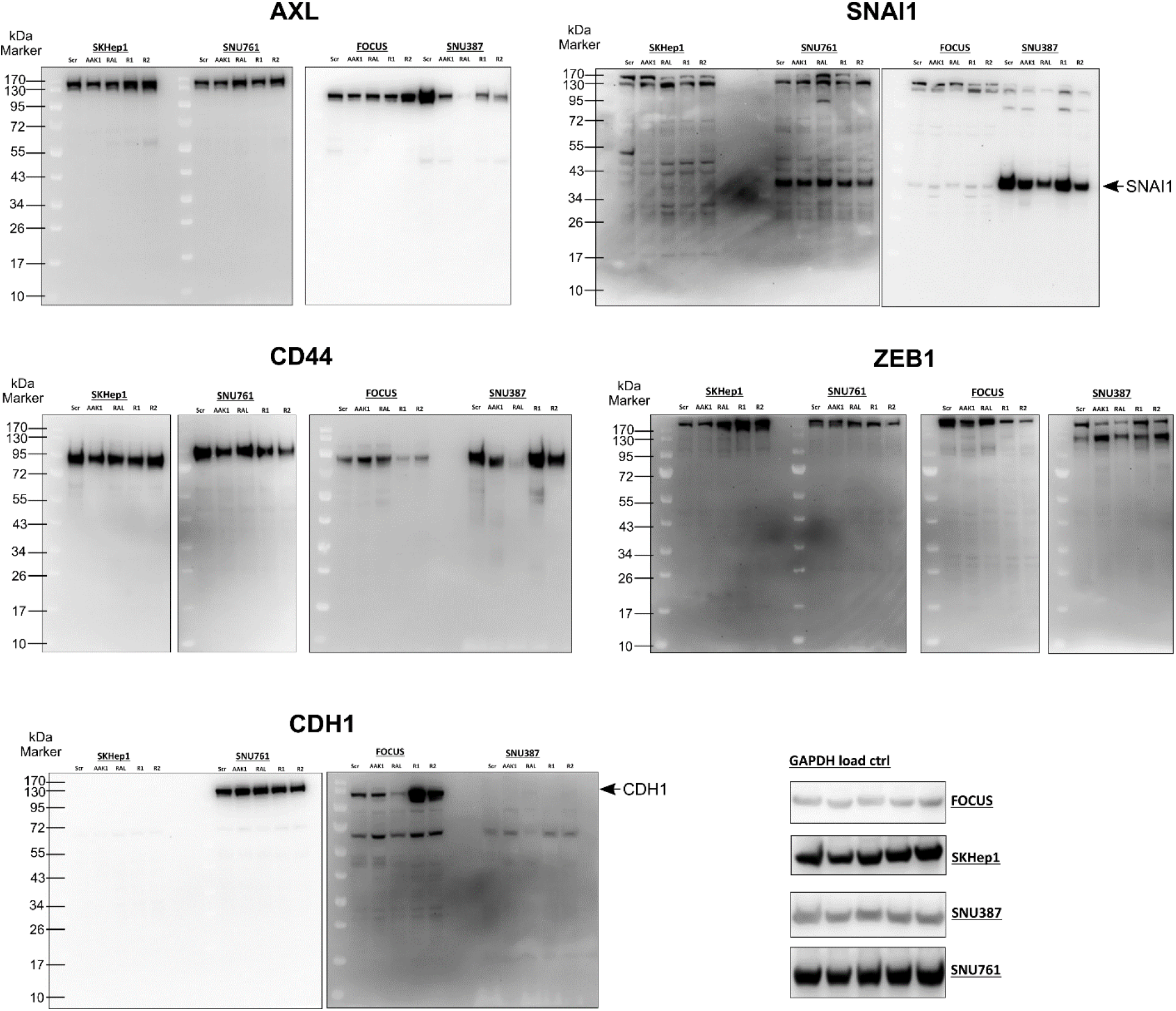
Immunoblot analysis of EMT marker expression in AAK1 complex RNAi cell lines.

## MATERIALS AND METHODS

### Cell lines and tissue culture conditions

C3A, SNU398, Hep3B2.1-7, U2-OS, SK-N-SH, SH-SY5Y, JeKo-1, HeLa, Jurkat, A-172, K562, and SNU449 cell lines were purchased from the American Type Culture Collection (ATCC). SNU761 and SNU886 were purchased from the Korean Cell Line Bank (KCLB). JHH6 and HuH-7 cells were purchased form the JRCB Cell Bank. FOCUS WT cells were obtained from the Laboratory of J. Wands, Brown University ^55^. FOCUS *AXL* RNAi cells were obtained from Dr. Taranjit Gujral of the Fred Hutchinson Cancer Research Institute, Seattle. All cells were grown at 37°C under 5% CO2, 95% ambient atmosphere. Fifteen cryo-frozen cell stocks were generated from the original vial from the cell bank (Passage 3). Experiments were performed with cells at <10 passages from the original vial. All cell media used were supplemented with 100x penicillin-streptomycin-glutamine (Thermo Fisher Scientific, Waltham, MA). FOCUS and HuH-7 cells were grown in Dulbecco’s minimum essential medium (DMEM) supplemented with 10% FBS (VWR Life Science, Seradigm). C3A, SNU398, Hep3B2.1-7, U2-OS, SK-N-SH, SH-SY5Y, JeKo-1, HeLa, Jurkat, A-172, K562, and SNU449 lines were grown in the ATCC-recommended medium. JHH6 cells were grown in William’s E medium, and SNU761 and SNU886 lines in RPMI 1640 medium all supplemented with 10% FBS. Cells were harvested when reaching 90% of confluency or a density of 1×10^6^ cells/ml.

### RNAi knockdown experiments

Three shRNA sequences each targeting AAK1, RALBP1, REPS1, and REPS2 were obtained from The RNAi Consortium (TRC) of the Broad Institute web portal (https://www.broadinstitute.org/rnai-consortium/rnai-consortium-shrna-library, ID Numbers: TRCN0000001943, TRCN0000199939, TRCN0000082348, TRCN0000053363, TRCN0000423162, TRCN0000436095, TRCN0000423057, TRCN0000428939, TRCN0000056210, TRCN0000305689, TRCN0000047918, TRCN0000047920) and cloned into the lentiviral pLKO.1 vector (Addgene, Watertown, MA) as previously described ^56^. Lentiviral particles were produced from individual pLKO.1 vectors, the pMD2.G plasmid (envelop), and the pCMVR8.74 plasmid (packaging) according to the manufacturer’s instructions (Addgene). Virus particle-containing cell culture supernatants were sterile filtered over 0.22 µM PES syringe filters (Millex-GP, Sigma Millipore, Burlington, MA), mixed 1:1 with fresh growth medium, 8 µg/mL polybrene was added and the mixture added to target cells (70-80% confluency). Cells were incubated for 24h, the medium exchanged, and stable cell lines selected using puromycin (FOCUS: 4 µg/mL; SNU387 and SKHep1: 6 µg/mL; SNU761: 8 µg/mL) for 7-14 days. Puromycin-resistant cells were maintained in growth medium containing half of the selection concentration of puromycin. Target knockdown was validated using qPCR and immunoblotting, and the stable cell lines with the highest knockdown among the three shRNAs used for each target were chosen to perform all downstream experiments.

### Immunoblot analysis and antibodies

Antibodies used were anti-E-cadherin (24E10, Cell Signaling Technology, CST, Cat # 3195), anti-AXL (C89E7, CST, Cat # 8661), anti-Snail (C15D3, CST, Cat # 3879), anti-ZEB1 (E2G6Y, CST, Cat # 70512), anti-CD44 (E7K2Y, CST, Cat # 37259), anti-GAPDH HRP conjugate (D16H11, CST, Cat # 8884), anti-AAK1 (E8M3P, CST, Cat # 61527), anti-RALBP1 (D87H8, CST, Cat # 5739), and anti-REPS1 (D6F4, CST, Cat # 6404). Cell lysis and immunoblotting experiments were performed using standard procedures. Briefly, cells were rinsed twice with ice-cold phosphate buffered saline (PBS), lysed in modified RIPA buffer V1 (50 mM Tris-HCl, 150 mM NaCl, 1% NP-40 (v/v), 0.25% Na-deoxycholate (w/v), 1 mM EDTA, 10 mM NaF, 5% glycerol (v/v), pH 7.8) supplemented with HALT protease inhibitor (100x, Thermo Fisher Scientific, Waltham, MA), and lysates clarified by centrifugation at 21,000 rcf for 20 minutes at 4°C. Protein concentration was quantified using the Piece 660 nm Protein Assay Reagent (Pierce, Rockford, IL). Lysates were mixed with NuPAGE LDS Sample Buffer (4X, Thermo Fisher Scientific) containing 50 mM DTT and heated for 5 min at 95°C. 20 µg of protein were separated on Bolt 4-12% Bis-Tris Protein Gels (Thermo Fisher Scientific) and electro-transferred onto nitrocellulose membranes. The buffer used for blocking and antibody incubation was 5% BSA in TBS-T (50 mM NaCl, 150 mM Tris-HCl, 1% Tween-20). Membranes were incubated with goat anti-rabbit HRP conjugate, and bands visualized using the Clarity Western ECL Substrate (Bio-Rad, Hercules, CA) and the Fluor Chem E imaging system (Protein Simple, San Jose, CA).

### Quantitative Real-Time PCR (qPCR) Analysis of mRNA Expression

shRNA-mediated knockdown was validated by quantifying mRNA expression levels using quantitative real-time PCR (qPCR). Briefly, cells were cultured on 35 mm dishes until reaching 80-90% confluency and total RNA was isolated using the TRIzol reagent according to manufacturer’s instructions (Thermo Fisher Scientific). mRNA quality was controlled by running 1% agarose gels and checking the presence of sharp, clear 28S and 18S rRNA bands. 0.5 µg of total RNA was used to generate first-strand cDNA using the Protoscript II First Strand cDNA Synthesis Kit (New England Biolabs, Ipswich, MA). The resulting cDNA was subjected to qPCR using human gene-specific primers for AAK1, RALBP1, REPS1, and REPS2, and two housekeeping genes, i.e., PSMB2 and RAB7A. The qPCR reaction was performed using QuantStudio 5 Real-Time PCR System (Applied Biosystems, Thermo Fisher Scientific) using the following program:

| Step | Temp (°C) | Time (mm:ss) | Cycles |
| --- | --- | --- | --- |
| Hold (enzyme active) | 50, 95 | 02:00, 10:00 | 1 |
| PCR (denature, anneal, extend) | 95, 50, 60 | 00:15, 00:15, 01:00 | 50 |
| Dissociation/melting curve | 95, 60, 95 | 00:15, 01:00, 00:15 | 5 |

The mRNA levels of each gene were normalized relative to the mean levels of the two housekeeping genes and compared with the data obtained from cell lines carrying a stably incorporated scramble shRNA using the 2-ΔΔCt method. According to this method, the normalized level of a mRNA, X, is determined using **Equation 1**:

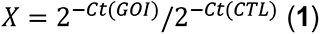

where Ct is the threshold cycle (the number of the cycle at which an increase in reporter fluorescence above a baseline signal is detected), GOI refers to the gene of interest, and CTL refers to a control housekeeping gene. This method assumes that Ct is inversely proportional to the initial concentration of mRNA and that the amount of product doubles with every cycle.

### Inhibitor treatment of RNAi lines and growth inhibition assay

1800 cells/well were seeded onto white flat bottom half area 96-well plates (Greiner Bio-One, Kremsmünster, AT) in 50 µl of growth medium and allowed to attach in an incubator for 24 h. Then the drugs in DMSO and/or DMSO vehicle controls as 11X solutions in growth medium were added to a total volume of 55 µl and 0.1% DMSO final. The cells were grown in an incubator for another 72 h. Then, 55 µl of CellTiter-Glo 2.0 (Promega, Madison, WI) reagent/well were added according to the manufacturer’s instructions and luminescence was quantified with a SpectraMax 190 plate reader (Molecular Devices, San Jose, CA). The CHEK1/2 inhibitors AZD7762 (Selleckchem, Houston, TX) and CHIR-124 (ApexBio, Houston, TX) were applied at 8 different concentrations between 10 µM and 4.6 nM (3-fold dilution steps). Experiments were performed in four biological replicates. Growth inhibition curves were fitted using the GraphPad Prism software package (V5.0a) with a least-squares nonlinear regression model for curve fitting (One site -Fit logIC50 function).

### Preparation of optimized kinobead mixture

The seven kinobead affinity reagents used were synthesized in-house as described previously ^19,21,57^. For optimal coverage of the human kinome an optimized mixture of the seven kinobead reagents was prepared as previously described ^19^. Briefly, 1 ml of reagent **1**, 0.5 ml of reagents **2**, **3** and **7**, respectively, and 0.25 ml of reagents **4**, **5** and **6**, respectively, were mixed to yield 3.25 ml of the complete kinobead mixture. All reagents were a 50% slurry in 20% aq. ethanol.

### Kinase affinity enrichment, KI competition and on-bead digestion

Kinase affinity enrichment, KI competition, and on-bead digestion was performed as previously described ^21,23,58^. Briefly, to 150 µL of cell lysate (5 mg protein per mL) in modified RIPA buffer V1 (50 mM Tris-HCl, 150 mM NaCl, 1% NP-40 (v/v), 0.25% Na-deoxycholate (w/v), 1 mM EDTA, 10 mM NaF, 5% glycerol (v/v), pH 7.8) containing HALT protease inhibitor cocktail (100x, Thermo Fisher Scientific, Waltham, MA) and phosphatase inhibitor cocktail II and III (100x, Sigma-Aldrich, St Louis, MO) 1.5 µL DMSO (vehicle control) or the corresponding inhibitor solution in DMSO (competition) were added to a final concentration 1% DMSO. The lysate was vortexed at intermediate speed intermittently every 5 min for 20 min while being kept on ice. Meanwhile, 40 µl of a 50% slurry of the in-house-made, optimized kinobead mixture in 20% aq. ethanol were prepared for each pulldown experiment. The beads were washed twice with 400 µl modified RIPA buffer and lysates containing DMSO, or inhibitor were added. The mixture was incubated on a tube rotator for 3h at 4°C and then the beads were spun down rapidly at 2000 rcf on a benchtop centrifuge (5s). After removal of the supernatant, the beads were rapidly washed twice with 400 µl of ice-cold mod. RIPA buffer and three times with 400 µl ice-cold tris-buffered saline (TBS, 50 mM tris, 150 mM NaCl, pH 7.8) to remove detergents. 100 µl of freshly prepared denaturing buffer (8M urea, 100 mM Tris, pH 8.5) containing 5 mM tris(2-carboxyethyl)phosphine hydrochloride (TCEP*HCl) and 10 mM chloroacetamide (CAM), were added and the slurry agitated on a thermomixer at 37°C and 1400 rpm for 30 min. The mixture was diluted 2-fold with 100 mM triethylamine bicarbonate (TEAB), the pH adjusted to 8-9 by addition 1 N aq. NaOH; 2 µg LysC were added, and the mixture agitated on a thermomixer at 1400 rpm at 37°C for 2 h. Then, the mixture was diluted another 2-fold with 100 mM TEAB, 2 µg MS-grade trypsin (Thermo Fisher Scientific, Waltham, MA) were added, and the mixture agitated on a thermomixer at 1400 rpm at 37°C overnight. Then, 6 µL of formic acid (FA) were added (1.5% FA final) to adjust to pH 3 and peptides were extracted ad desalted using C18 StageTips according to the published protocol ^59^. For kinobead/LC-MS profiling of RNAi cell lines the same protocol was applied except that lysates were not preincubated with DMSO or inhibitor. The following inhibitors were used for competition experiments at the given final concentrations: GSK-690693 (10 µM, MedChemExpress, MCE, Monmouth Junction, NJ), Miliciclib (10 µM, MCE), Rebastinib (10 µM, MCE), AT9283 (10 µM, MCE), TAK-901 (10 µM, MCE), RGB-286638 (10 µM, MCE), Flavopiridol*HCl (10 µM, MCE), PF-562271 besylate (10 µM, MCE), Dabrafenib mesylate (10 µM, MCE), OTSSP167*HCl (10 µM, MCE), CYC-116 (10 µM, MCE), Silmitasertib (10 µM, MCE), SB1317 (10 µM, MCE), XL228 (10 µM, MCE), Sapanisertib (10 µM, MCE), PF-3758309 (10 µM, ApexBio), Staurosporine (1 µM, LC Labs, Woburn, MA), AZD-7762 (10 µM Selleckchem), Bosutinib (10 µM, Selleckchem), Dasatinib (10 µM, Selleckchem), Linsitinib (10 µM, ApexBio).

### Ki-CCA with formaldehyde-mediated protein crosslinking

Ki-CCA with formaldehyde-mediated protein crosslinking was performed as described in “Kinase affinity enrichment, KI competition, and on-bead digestion” above with the following modifications. Briefly, to 150 µL of cell lysate (5 mg protein per mL) in modified RIPA buffer V2 (50 mM HEPES, 150 mM NaCl, 1% NP-40 (v/v), 0.25% Na-deoxycholate (w/v), 1 mM EDTA, 10 mM NaF, 5% glycerol (v/v), pH 7.8) containing HALT protease inhibitor cocktail (100x, Thermo Fisher Scientific, Waltham, MA) and phosphatase inhibitor cocktail II and III (100x, Sigma-Aldrich, St Louis, MO) 1.5 µL DMSO (vehicle control) or the corresponding inhibitor solution in DMSO (competition) were added to a final concentration 1% DMSO. The lysate was vortexed at intermediate speed intermittently every 5 min for 20 min while being kept on ice. The mixture was added to the kinobeads and incubated on a tube rotator for 3h at 4°C and then 4 µL of 37 wt% aq. Formaldehyde solution was added (1% concentration final). The mixture was incubated on a tube rotator for an additional 30 min at 4°C and then the beads were spun down rapidly at 2000 rcf on a benchtop centrifuge (5s). After removal of the supernatant, the beads were rapidly washed twice with 400 µl of ice-cold mod. RIPA buffer V2 and three times with 400 µl ice-cold HEPES-buffered saline (HBS, 50 mM HEPES, 150 mM NaCl, pH 7.8) to remove detergents. 100 µl of the denaturing buffer (6M Gdn*HCl, 100 mM Tris-HCl, pH 8.5) containing 5 mM tris(2-carboxyethyl)phosphine hydrochloride (TCEP*HCl) and 10 mM chloroacetamide (CAM), were added and the slurry agitated on a thermomixer at 70°C and 1400 rpm for 30 min to reverse crosslinking. The mixture was then subjected to the same digestion protocol and downstream handling as described above.

### Co-Immunoprecipitation of AAK1 Complex Components

200 µl of FOCUS cell lysate in modified RIPA buffer V1 (5 mg/mL protein) containing protease and phosphatase inhibitors (see ‘Kinase affinity enrichment, KI competition and on-bead digestion’) were incubated with antibodies against AAK1, RALBP1, REPS1 (see ‘Immunoblot analysis and antibodies’), or GFP (control, D5.1, CST, Cat # 2956) at the manufacturer’s recommended concentrations, respectively, and agitated overnight on a tube rotator at 4°C. The next day, Pierce Protein A Agarose (Thermo Fisher Scientific) was washed twice with ice-cold modified RIPA buffer V1 and 30 µL aliquots of a 50% bead slurry were added to each lysate/antibody mixture. The slurry was agitated for 3h on a tube rotator at 4°C and then the beads were washed two times with ice-cold modified RIPA buffer and three times with TBS. Proteins were reduced, alkylated, and eluted with 100 µL denaturing buffer (8M urea, 2mM TCEP, 4mM CAM, 100 mM Tris pH 7.8) for 30 min. The supernatant was transferred to a fresh tube and diluted two-fold with 100 mM TEAB, and the pH adjusted to 8.5 using 1N NaOH solution. 2 µg of Lys-C (Wako Chemicals) were added, and samples incubated at 1,400 rpm at 37°C for 2 h on a thermo mixer. Then, sampled were diluted another two-fold with 100 mM TEAB and 2 μg of MS-grade trypsin (Thermo Fisher Scientific) were added. The mixture agitated on a thermomixer at 1,400 rpm at 37°C overnight. The digests were acidified with formic acid to pH <3 (1.5% FA final) and desalted using C18 StageTips according to the published protocol ^59^. Co-IP-MS experiments were performed in three biological replicates per antibody.

### nanoLC-MS/MS analyses

LC-MS analyses were performed as described previously with the following minor modifications ^19,21^. Peptide samples were separated on an EASY-nLC 1200 System (Thermo Fisher Scientific) using 20 cm long fused silica capillary columns (100 µm ID) packed with 3 μm 120 Å reversed phase C18 beads (Dr. Maisch, Ammerbuch, DE). The LC gradient was 120 min long with 5−35% B at 300 nL/min. LC solvent A was 0.1% (v/v) aq. acetic acid and LC solvent B was 20% 0.1% (v/v) acetic acid, 80% acetonitrile. MS data was collected with a Thermo Fisher Scientific Orbitrap Fusion Lumos. Data-dependent analysis was applied using Top15 selection with CID fragmentation.

### Computation of MS raw files

Data .raw files were analyzed by MaxQuant/Andromeda ^60^ version 1.5.2.8 using protein, peptide and site FDRs of 0.01 and a score minimum of 40 for modified peptides, 0 for unmodified peptides; delta score minimum of 17 for modified peptides, 0 for unmodified peptides. MS/MS spectra were searched against the UniProt human database (updated July 22nd, 2015). MaxQuant search parameters: Variable modifications included Oxidation (M) and Phospho (S/T/Y). Carbamidomethyl (C) was a fixed modification. Max. missed cleavages was 2, enzyme was Trypsin/P and max. charge was 7. The MaxQuant “match between runs” feature was enabled. The initial search tolerance for FTMS scans was 20 ppm and 0.5 Da for ITMS MS/MS scans.

### MaxQuant output data processing

MaxQuant output files were processed, statistically analyzed and clustered using the Perseus software package v1.5.6.0 ^61^. Human gene ontology (GO) terms (GOBP, GOCC and GOMF) were loaded from the ‘Perseus Annotations’ file downloaded on 01.08.2017. Expression columns (protein and phosphopeptide intensities) were log2 transformed and normalized by subtracting the median log2 expression value from each expression value of the corresponding data column. Potential contaminants, reverse hits and proteins only identified by site were removed. Reproducibility between LC-MS/MS experiments were analyzed by column correlation (Pearson’s r) and replicates with a variation of r > 0.25 compared to the mean r-values of all replicates of the same experiment (cell line or knockdown experiment) were considered outliers and excluded from the analyses. Data imputation was performed using a modeled distribution of MS intensity values downshifted by 1.8 and having a width of 0.2.

### Kinobead competition correlation analysis (Ki-CCA)

For each cell line tested, 21 KIP competition experiments and one DMSO control experiment were performed in biological duplicate, resulting in 44 kinobead pulldown and LC-MS experiments per condition/cell line. We defined a kinase or non-kinase protein as competed if it showed a log2 MS intensity ratio of ≥ 0.75 and passed a two-sample t-test p < 0.1 with at least one of the 21 KIs used, i.e., comparing the two DMSO control experiments and two corresponding KI competition experiments. We then extracted the log2-transformed, normalized, and imputed MS intensity values of all competed kinases and non-kinases and correlated all kinases with all non-kinases for individual cell lines using Pearson moment correlation (n = 44). Then, kinases that show similar competition behavior were combined into groups (***see below***), and the maximum r-value of members retained for that group. Finally, the kinase group which showed the highest r-value for each non-kinase protein was determined, presenting the most probably kinase group – protein interaction among all possible interaction for each cell line.

### Determining kinase groups with similar KIP binding profiles

Like Ki-CCA correlation analysis, kinase MS Intensity values were correlated with one another for each cell lines tested to identify kinases with very similar competitor binding profiles. Thus, kinases that show an r-value > 0.9 in at least 7 of 21 tested cell line were combined into kinase groups, defining that the interactors of kinases in these groups cannot be distinguished using Ki-CCA. Examples are PRKAA1 and 2, AAK1 and BMPK, as well as STK24 and STK26 (see Table S2, Tab ‘Kinase Groups’)

### Plotting STRING Interaction Networks

PPI networks were plotted using the STRING web application version 11.5 with the following settings: Edges were scaled with confidence, and only ‘physical subnetwork’ interactions were considered, i.e., only considering text mining, experiments, and databases ^62^.

### KS-test analysis and Receiver Operating Characteristic (ROC) plots

Combined PPIs from BioGrid, BioPlex, and Buljan et. al. 2020 were used to populate the ‘known’ PPIs in our dataset and used as the binary classifier. The Pearson’s r is used as the discrimination variable. KS plots were generated with the ‘ROCit’ package in R. ROC p-values were determined with the ‘verification’ package in R.

### Kinome dendrograms

Kinome dendrograms were prepared using the KinMap web application (http://kinhub.org/kinmap/)^63^.

